# A Panoramic View of Cell Population Dynamics in Mammalian Aging

**DOI:** 10.1101/2024.03.01.583001

**Authors:** Zehao Zhang, Chloe Schaefer, Weirong Jiang, Ziyu Lu, Jasper Lee, Andras Sziraki, Abdulraouf Abdulraouf, Brittney Wick, Maximilian Haeussler, Zhuoyan Li, Gesmira Molla, Rahul Satija, Wei Zhou, Junyue Cao

## Abstract

To elucidate the aging-associated cellular population dynamics throughout the body, here we present PanSci, a single-cell transcriptome atlas profiling over 20 million cells from 623 mouse tissue samples, encompassing a range of organs across different life stages, sexes, and genotypes. This comprehensive dataset allowed us to identify more than 3,000 unique cellular states and catalog over 200 distinct aging-associated cell populations experiencing significant depletion or expansion. Our panoramic analysis uncovered temporally structured, organ- and lineage-specific shifts of cellular dynamics during lifespan progression. Moreover, we investigated aging-associated alterations in immune cell populations, revealing both widespread shifts and organ-specific changes. We further explored the regulatory roles of the immune system on aging and pinpointed specific age-related cell population expansions that are lymphocyte-dependent. The breadth and depth of our ‘cell-omics’ methodology not only enhance our comprehension of cellular aging but also lay the groundwork for exploring the complex regulatory networks among varied cell types in the context of aging and aging-associated diseases.

**One Sentence Summary:** PanSci, a single-cell transcriptome atlas of over 20 million cells throughout the mouse lifespan, unveils the temporal architecture of aging-associated cellular population dynamics, organ-specific immune cell shifts, and the lymphocyte’s role in organismal aging.

## Main Text

The adage ‘A chain is only as strong as its weakest link’ aptly applies to the aging process. Within the diverse cellular landscape of different organs, certain cell types exhibit profound alterations in their states or populations as we age(*1–3*). These changes not only impact the overall function of the organism, but play a critical role in the onset of age-associated diseases(*4*). Therefore, cataloging these vulnerable cell types is critical for unraveling the cellular underpinnings of aging-related pathologies and for identifying potential interventions to counteract detrimental age-related changes in cell populations.

Nevertheless, a comprehensive characterization of these aging-related cellular changes presents significant challenges. One primary barrier is the inherent heterogeneity within cell populations, potentially obscuring less common yet crucial cell types involved in aging. While advancements in single-cell genomic profiling offer a powerful route in characterizing cell state heterogeneity(*5–8*), current studies are limited by the throughput of single-cell techniques and thus mainly focus on the abundant cell types, neglecting the intricate dynamics of rare cell states or subtypes, as well as their variations across different individuals or conditions (*e.g.,* sexes, genotypes). In addition, large-scale single-cell studies that integrate multiple datasets, often profiled using varied methodologies and by different laboratories, face the challenge of batch effects that can hinder the identification of rare cell types and complicate comparisons of broadly distributed cell types, such as immune or endothelial cells, across different tissues(*9–11*).

To achieve a comprehensive characterization of aging-associated cell population changes, here we present *PanSci*, a panoramic view of mouse aging, by examining the transcriptional states of over twenty million cells across mammalian organs sourced from 623 diverse tissue samples (**Fig. 1A-B, fig. S1-2**). These samples were collected from a cohort of individuals across various ages, sexes, and genotypes (**fig. S2A-C, table S1**). Specifically, we included eight sex-balanced wild-type C57BL/6 mice across three age groups (6-month, 12-month, and 23-month). Moreover, we profiled both wild-type and two immuno-deficient genotypes, *B6.129S7-Rag1tm1Mom/J*(*12*) and *B6.Cg-Prkdcscid/SzJ*(*13*), at 3-month and 16-month stages, with four sex-balanced replicates each. (**Fig. 1A, upper**). These mutant strains, characterized by lymphocyte deficiency, could provide insight into the regulatory role of the immune system in the aging process of other solid organs. In addition, varied time intervals and genotypes in our dataset allow for rigorous cross-validation of the observed aging-associated cell population changes.

**Figure 1:**
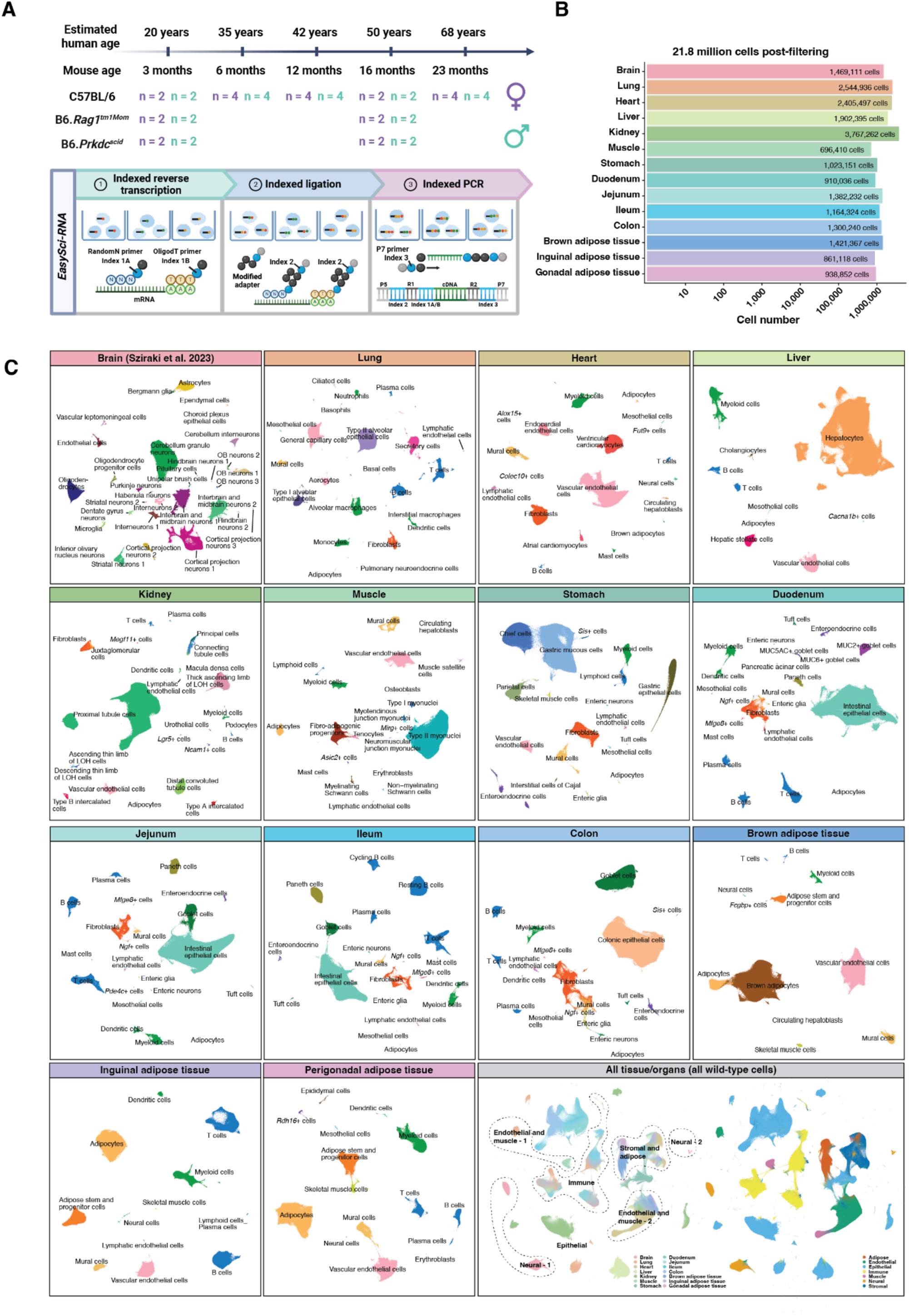
Overview of experimental design and main cell type annotation across mammalian organs. (**A**) Upper: Schematic representation of the sample collection process detailing the various ages, sexes, and genotypes (including wild-type and immuno-deficient mice) used in the study. Lower: Flowchart illustrating the experimental procedures of single-cell RNA sequencing by combinatorial indexing through *EasySci*. (**B**) Logarithmic scale bar plot depicting the number of high-quality cells profiled from each organ or tissue, post-quality filtering. (**C**) UMAP plots displaying the cellular heterogeneity of each organ/tissue, with cells color-coded by identified main cell types. Brain cell types were retrieved from (*14*). An aggregated UMAP plot of the entire dataset (comprising only wild-type cells, without batch correction) is also shown (right corner), with cells distinguished by organ/tissue origin and lineage. LOH, loop of Henle.

The single-cell datasets are generated with *EasySci*(*14*), an optimized single-cell combinatorial indexing method(*15–18*) for organismal cell population analyses. A noteworthy aspect of *EasySci* lies in its full gene body coverage of transcripts, scalability, and cost-effectiveness, facilitating the profiling of over 20 million cells by a single operator. Notably, we extensively optimized the cell lysis conditions to efficiently extract nuclei from frozen tissues across diverse mammalian organs, effectively reducing the batch effects commonly associated with conventional tissue digestion and cell isolation approaches (**fig. S1**). Post extraction, the nuclei underwent fluorescence-activated cell sorting, followed by barcoding via indexed reverse transcription, ligation, and PCR stages in *EasySci* (**Fig. 1A, lower**). The final libraries were sequenced through twenty-five S4 runs with the Illumina NovaSeq 6000 system, yielding over 200 billion raw reads. This sequencing depth (∼8,950 reads per cell), aligns with our prior single-cell studies in capturing rare cell states in mammalian development and brain aging (*14*, *18*, *19*). After filtering low-quality cells and doublets, we recovered 21,786,931 single-nucleus gene expression profiles (including the 1,469,111 brain cells profiled in(*14*)) (**fig. S2G**). An average of 1,601 unique transcripts (UMIs) was detected per cell (median = 1,040 UMIs) (**fig. S2D**), and an average of 1,562,909 cells was profiled per organ (**Fig. 1B**; maximum, 3,767,262 cells from kidney; minimum, 696,410 cells from muscle).

We adopted a two-step approach, akin to our prior work(*14*), to identify heterogeneous cellular states across various organs. Employing UMAP visualization and Leiden clustering(*20*), we first analyzed single-cell gene expression profiles for each organ separately. A total of 239 organ-specific main cell types were characterized across different organs (except the 31 brain cell types identified in (*14*)) (**Fig. 1C, fig.S3A, table S2**). Each cell type was identified across multiple individuals (a median of 48 samples per cell type, **fig. S3B**), represented by a median of 15,922 cells, ranging from 2,465,275 cells (*i.e.,* proximal tubule cells in the kidney) to only 6 cells (*i.e.,* osteoblasts in muscle) (**fig. S3C**). On average, 56 unique marker genes were identified per cell type. The marker genes were defined by a minimum fivefold difference in expression between the top-ranked and second-ranked cell types and a minimum expression (transcripts per million) of 50 in the targeted cell type (**table S3**). The identity of these cell types was confirmed by cell-type-specific gene markers from published single-cell datasets (*10*, *21–33*) (**fig. S4-6**). Notably, the scalability of our platform has effectively minimized the batch effects that arise during the integration of single-cell datasets generated in multiple laboratories in conventional consortium-level studies (*6*, *34*). Taking the muscle as an example, cells of the same type (*e.g.,* Type II myonuclei) from different individuals are clustered together in the UMAP space without batch correction (**fig. S7A-B**). A subsequent validation, incorporating cells from various organs, confirmed that broadly distributed cell types, such as immune and endothelial cells, were clustered together in the UMAP space (**Fig. 1C**).

As a second step toward a more detailed characterization of cellular heterogeneity, we took each main cell type for sub-clustering analysis by integrating both gene and exonic counts per cell(*14*). This is based on a unique feature of the *EasySci* approach that integrates both indexed oligo-dT primers and random primers during reverse transcription, ensuring full gene body coverage and simultaneous recovery of non-polyA transcripts. Similar to our previous study(*14*), the combined information remarkably increased the clustering resolution (**fig. S7C**). Beyond the 359 sub-clusters we identified before in the brain(*14*), we detected 3,925 sub-clusters across organs, with each observed in multiple individuals (a median of 45 per sub-cluster), represented by a median of 1,035 cells (**fig. S3D-F)**. Over 90% of these sub-clusters (3,535 out of 3,925) can be distinguished by unique gene markers per the above-mentioned criteria (**table S4**). To validate the unique transcriptomic signatures of these sub-clusters, we harnessed 80% of our single-cell gene expression dataset to train a support vector machine classifier for sub-cluster annotation. This classifier, upon application to the residual dataset, recognized most sub-clusters compared with permutation controls (**fig. S7D-G**).

By incorporating sex-balanced replicates into each group, our dataset provides an in-depth view of the sex-specific effects on heterogeneous cellular states across various organs. Taking the liver as an example, we observed distinct separations in specific cell populations between females and males in hepatocytes (**fig. S8A-B**), consistent with previous research characterizing the liver as a “sexually dimorphic organ”(*35*, *36*). This distinction is in line with known sex-specific variations in metabolic functions, such as superior alcohol clearance and lipid metabolism capabilities in males and heightened cholesterol metabolism abilities in females(*36*). These sex-specific effects extend down to the sub-cluster level, as demonstrated by the identification of 73 sub-clusters displaying significant differential abundance between males and females across all age groups (**fig. S9**). Interestingly, our analysis not only reaffirmed the presence of conserved sexually dimorphic cell types in the liver and kidney (*37*), but also brought to light underreported cell types in other organs (**fig. S8C-F**). For instance, in the perigonadal adipose tissue (gWAT), we identified female-specific *Dlgap1+ Fgf10*+ and male-specific *Pde11a+ Rtl4*+ adipose stem and progenitor cells. In the stomach, we found female-specific *Grm8+ Entpd1*+ and male-specific *Slc35f3+ Rimbp2*+ Chief cells. These discoveries highlight the complexity of sex-specific cellular differences and pave the way for future in-depth studies.

### Temporally structured aging-associated cell population dynamic waves

To obtain a global view of aging-related cell population dynamics, we quantified cell-type-specific proportions in both main cell types and sub-clusters within individual replicates across various age groups, followed by differential abundance analyses (**Methods**). We detected 23 main cell types and 374 sub-clusters undergoing significant population changes in both age intervals: 3 vs. 16 months and 6 vs. 23 months (False discovery rate (FDR) of 0.05, with a minimum 2-fold difference between two-time points; **Fig. 2A, fig. S10**). These changes were confirmed by significant consistency between different sex (**Fig. 2B-E**). Reassuringly, most of these main cell types (21 out of 23) and sub-clusters (280 out of 374) demonstrated robust and consistent changes during both intervals, referred to as “aging-associated cell populations” in our subsequent analysis.

**Figure 2:**
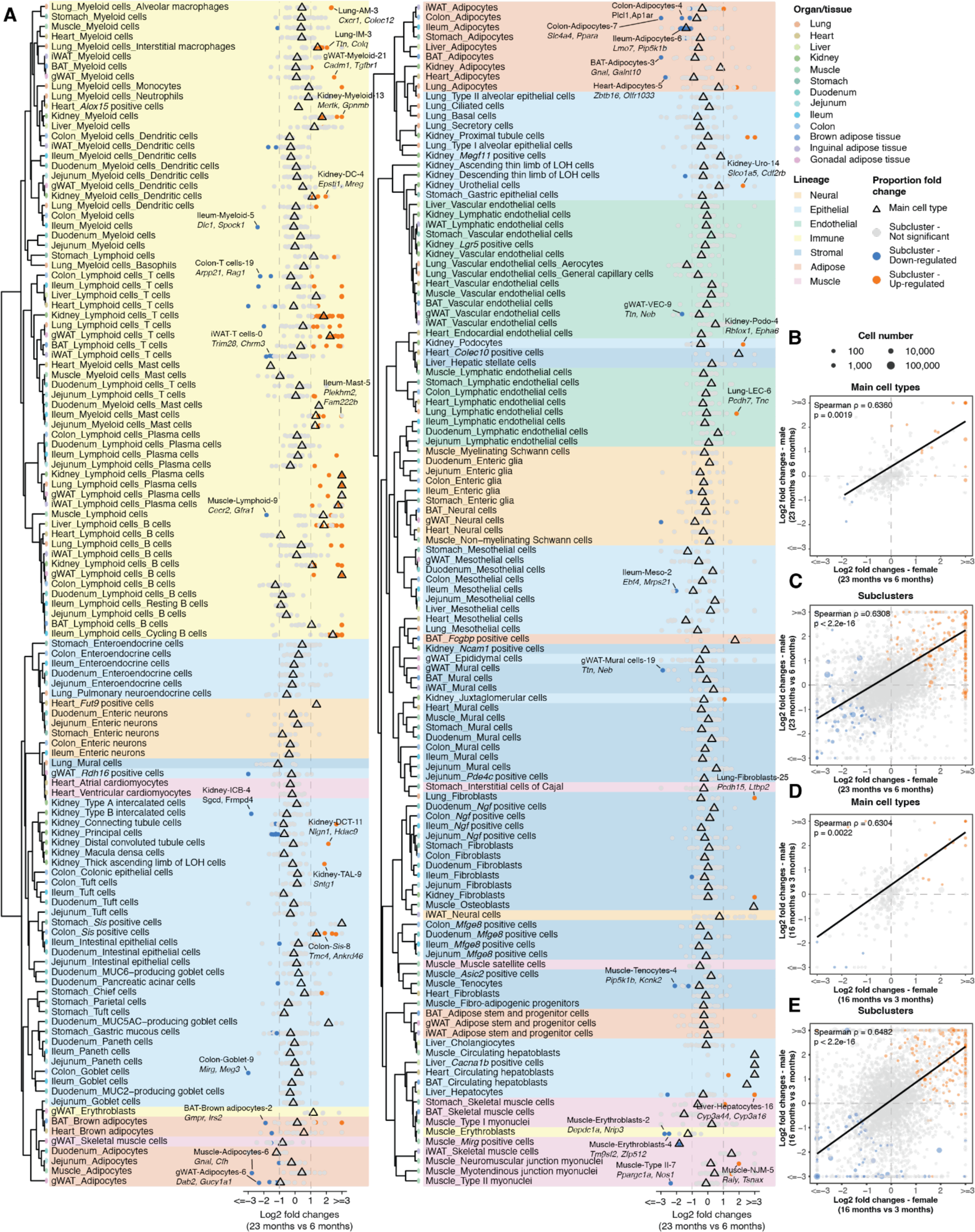
Identification of aging-associated cell population change across organs/tissues. (**A**) Dot plots illustrating cell-type-specific fractional changes (log-transformed fold change) between ages 6 and 23 months. Main cell types are represented by triangles and sub-clusters by dots, with key gene markers labeled for select sub-clusters. The dendrogram is derived from hierarchical clustering of gene expression correlations among main cell types. AM, alveolar macrophages; IM, interstitial macrophages; DC, dendritic cells; ICB, Type B intercalated cells; DCT, distal convoluted tubule cells; TAL, thick ascending limb of LOH cells; *Sis*, *Sis* positive cells; Uro, urothelial cells; VEC, vascular endothelial cells; Podo, podocytes; LEC, lymphatic endothelial cells; Meso, mesothelial cells; Type II, Type II myonuclei; NJM, neuromuscular junction myonuclei. (**B**-**E**) Correlation scatter plots (employing Spearman correlation) comparing fractional changes in main cell types (B, D) and sub-clusters (C, E) between female and male mice during two age intervals: 6 vs. 23 months (B, C) and 3 vs. 16 months (D, E), with a linear regression line. For all scatter plots, aging-associated cell types that are significantly changed in both age intervals are colored by the direction of changes.

These aging-associated cell populations exhibited unique dynamics across distinct life stages (**Fig. 3A**). For instance, we observed a significant age-associated expansion in immune cells, including lymphocytes and myeloid cells across multiple organs, aligning with findings from previous studies(*38*). Interestingly, certain age-associated cell types also exhibited a marked sexual dimorphism. For example, while we observed a decline in *Mirg*+ cells within the skeletal muscle across aging in both sexes, there is a sharper reduction in females due to a higher baseline level in youth (**Fig. 3B**). These cells correspond to a rare subset of muscle myonuclear populations, marked by an elevated expression of several lncRNAs—*Meg3, Rian, GM37899, and Mirg*—all stemming from the *Dlk1-Dio3* locus, containing mammalian’s largest miRNA mega-cluster(*39*)(**Fig. 3C-D**). The observed decline may be attributed to the aging-associated downregulation of miRNAs from *Dlk1-Dio3* locus, critical for mitochondrial biogenesis and reactive oxidative species protection, suggesting the diminished cellular resilience against aging-induced stress(*40*).

**Figure 3:**
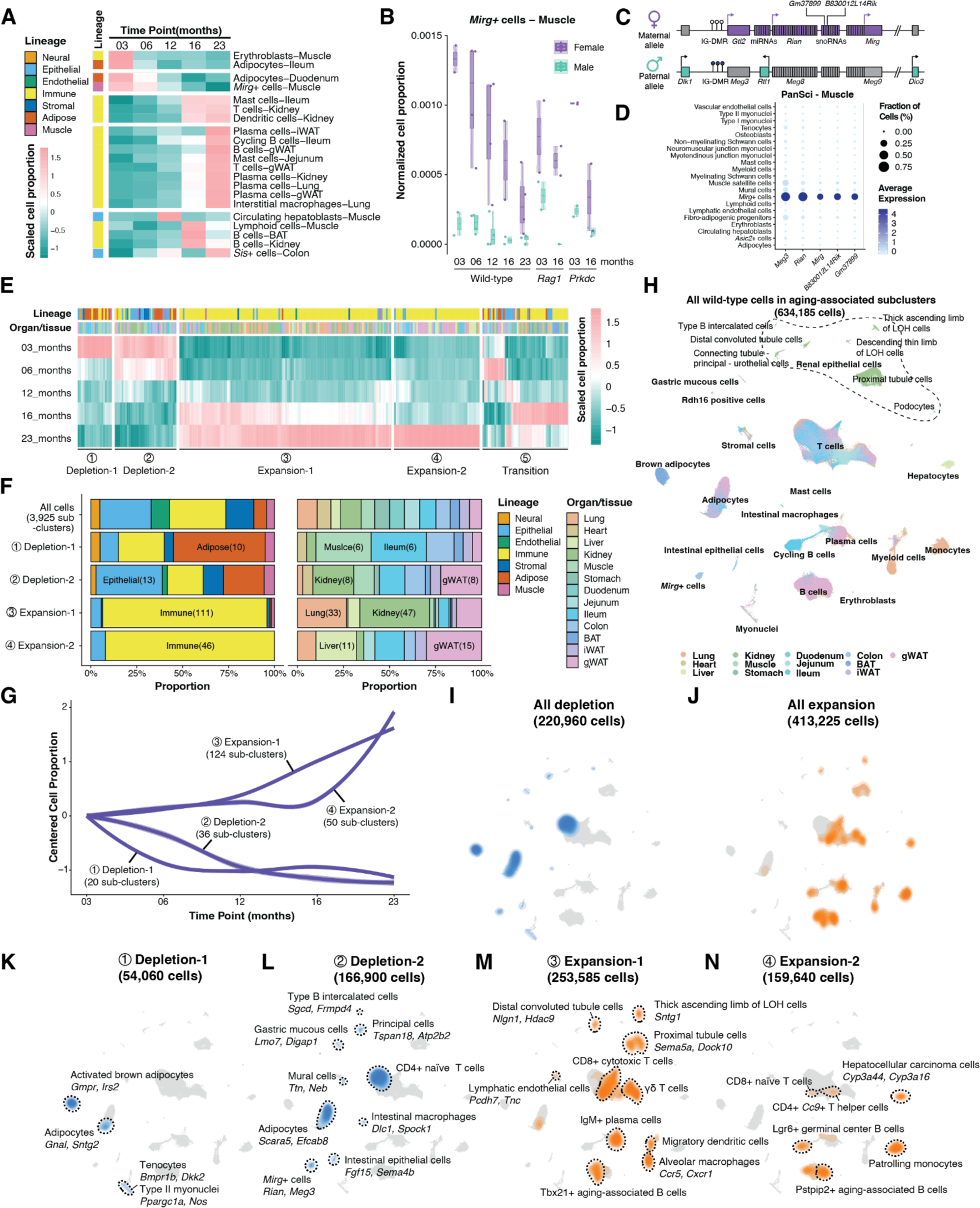
The temporal dynamics, tissue distribution, and molecular signatures of aging-associated cell populations. (**A**) Heatmap illustrating the fractional changes of aging-associated main cell types across five life stages. (**B**) Box plots depicting the fractional changes in muscle *Mirg*+ cells (lower) across the five life stages in wild-type and two time points in two lymphocyte-deficient mutants. Each dot represents a biological replicate. For all box plots: middle lines, median value; upper and lower box edges, first and third quartiles, respectively; whiskers, 1.5 times the interquartile range; and all individual data points are shown. (**C**) Schematic of the *Dlk1-Dio3* locus, highlighting *Mirg*+ cell marker genes. (**D**) Dot plot displaying marker gene expression in the PanSci-muscle dataset, with color indicating average expression and dot size showing the percentage of cells expressing each marker. (**E**) Heatmap of aging-associated sub-cluster fractions across five life stages, with hierarchical clustering identifying distinct depletion and expansion waves. (**F**) Stacked bar plots representing the proportions of aging-associated sub-clusters from different lineages and organs/tissues in each dynamic wave. (**G**) Line plot showing normalized cell proportion changes in each aging wave, with Loess regression lines centered at the initial age point. (**H**) UMAP visualizations of 634,185 wild-type cells from aging-associated sub-clusters, colored by organ/tissue. (**I**-**N**) Density plots showing the distribution of aging-associated sub-clusters from all depletion dynamic waves (I), all expansion dynamic waves (J), first depletion wave spanning 3 to 6 months (K), second depletion wave extending to 12 months (L), first expansion wave starting from 12 months (M), and second expansion wave from 16 months (N). Cells from non-immune lineage are annotated with enriched genes.

To delve deeper into the evolving landscape of cell populations across the lifespan, we next clustered all age-associated cell sub-clusters, based on their dynamics across five age stages. A substantial proportion of these sub-clusters underwent consistent alterations throughout life. Specifically, we identified 174 sub-clusters that expanded, 56 that depleted, and 50 with transient dynamics (**Fig. 3E**). Utilizing our multi-timepoint dataset, we discerned the varied pace at which different cellular states altered across cell lineages and organs (**Fig. 3F-G**). To further investigate these aging-associated cell populations and their unique molecular markers, we integrated all cells from consistently expanding or depleting sub-clusters for clustering and UMAP visualization (**Fig. 3H**). This analysis led to the characterization of distinct cellular dynamics at different life stages, accompanied by both organ- and lineage-specific cellular populations, as discussed below:

The initial two waves predominantly indicate cell loss (**Fig. 3I**). The first wave, spanning 3 to 6 months, is characterized by a decline in *Gmpr*+ activated brown adipocytes in BAT, accompanied by a reduction in *Ppargc1a*+ *Nos*+ type II myonuclei and *Bmpr1b*+ *Dkk2*+ tenocytes in muscle (**Fig. 3K**). The subsequent wave, extending from 6 to 12 months, is notable for the marked decrease in CD4+ naïve T cells across various organs and *Dlc1+ Spock1+* intestinal macrophages (**fig. S11D**). As a continuum of muscle degeneration, we witnessed a reduction of *Flt1*+ *Mecom*+ tenocytes from muscle and *Ttn+ Neb+* mural cells from gWAT. The age-related adipose decline is further observed in *Bmper*+ *Scara5*+ adipocytes from gWAT. Additionally, this period featured a decrease in functional epithelial cells across several organs, including *Mirg*+ cells from muscle and colon, *Fgr15*+ intestinal epithelial cells, *Lmo7+ Digap1+* gastric mucous cells, *Rdh16*+ cells from gWAT, and a range of epithelial cells in the kidney (*e.g., Sgcd+ Frmpd4*+ type B intercalated cells, *Zfp207*+ connecting tubule cells, and *Tspan18+* principal cells) (**Fig. 3L**).

The following two waves are featured with the cell expansion, primarily immune cells (**Fig. 3J**). The third wave, initiating from 12 months, is dominated by an expansive growth in the majority of T cell subtypes (*e.g.,* CD8+ *Gzmk*+ cytotoxic T cells, and Gamma-delta (γδ) T cells), IgM+ plasma cells, *Tbx21*+ aging-associated B cells, and various subtypes of cells from myeloid lineages (*e.g., Blnk+ Ccr5+* alveolar macrophages, *Cxcr1+ Fstl1+* alveolar macrophages, migratory dendritic cells, *Col14a1*+ macrophages from kidney, *Colq+* macrophages from intestine, *Mcpt1+ Mcpt2+* mucosal mast cells (**fig. S11E**)). Additionally, *Pcdh15+ Ltbp2+* lung fibroblasts, *Pcdh7+ Tnc+* lung lymphatic endothelial cells, *Raly+ Tsnax+* neuromuscular junction myonuclei, *Ampd1+ Cdc14a+* myotendinous junction myonuclei and certain epithelial cells in the kidney (*e.g., Sema5a+ Dock10+* proximal tubule cells, *Sntg1+* thick ascending limb of LOH cells, *Slco1a5+ Cdf2rb+* urothelial cells, *Rbfox1+ Epha6+* podocytes, *Prkca+ Nrxn3+* Juxtaglomerular cells, and *Nlgn1+ Hdac9+* distal convoluted tubule cells) significantly surged as well (**Fig. 3M**). The fourth wave, starting from 16 months, is characterized by a major expansion in immune populations, such as *Pstpip2*+ aging-associated B cells and patrolling monocytes (**Fig. 3N, fig. S11E**).

In summary, these sequential aging dynamics waves delineate a pattern where cellular depletion precedes expansion, with minimal temporal overlap, suggesting disparate mechanisms governing cell population dynamics at varying life stages. The initial dynamics, spanning from 3 to 12 months, are predominantly marked by the loss of cells in adipose, muscle, and epithelial lineages. Conversely, the latter stages, from 12 to 23 months, exhibit a significant expansion of immune cells. This progression aligns with prior research documenting sequential alterations of plasma protein profiles throughout the aging process(*41*), thereby reflecting the non-linear shifts of the internal milieu at different life stages.

### A global view of aging-associated changes in lymphocyte populations

Our pan-organ dataset provides a unique opportunity for systematically exploring organ-specific aging changes in broadly distributed cell types, particularly immune cells. To investigate the aging-related alteration of T cells and innate lymphoid cells (ILCs), we first isolated 957,975 cells representing these cell populations across all organs for clustering and UMAP visualization (**Fig. 4A**). A total of 18 cell clusters were recovered, each with highly cell-type-specific gene markers (**Fig. 4B**). While most cell clusters are prevalent across various organs, some immune cells exhibited organ-specific distribution (**Fig. 4C**). For instance, ILC-3 cells (Cluster 18), the central regulator of gut immunity(*42*), predominantly detected in the intestine. Similarly, *Prf+* natural killer cells (Cluster 13) – crucial for pathogenic immune response and maintaining pulmonary homeostasis(*43*) – were primarily found in the lung. Additionally, while CD8*+ Gzmb+* cytotoxic T cells (Cluster 8) were predominantly in the intestine, CD8*+ Gzmk+* cytotoxic T cells (Cluster 9) appeared more abundantly in other organs such as the kidney, lung, and adipose tissue, aligning with the prior report(*44*).

**Figure 4:**
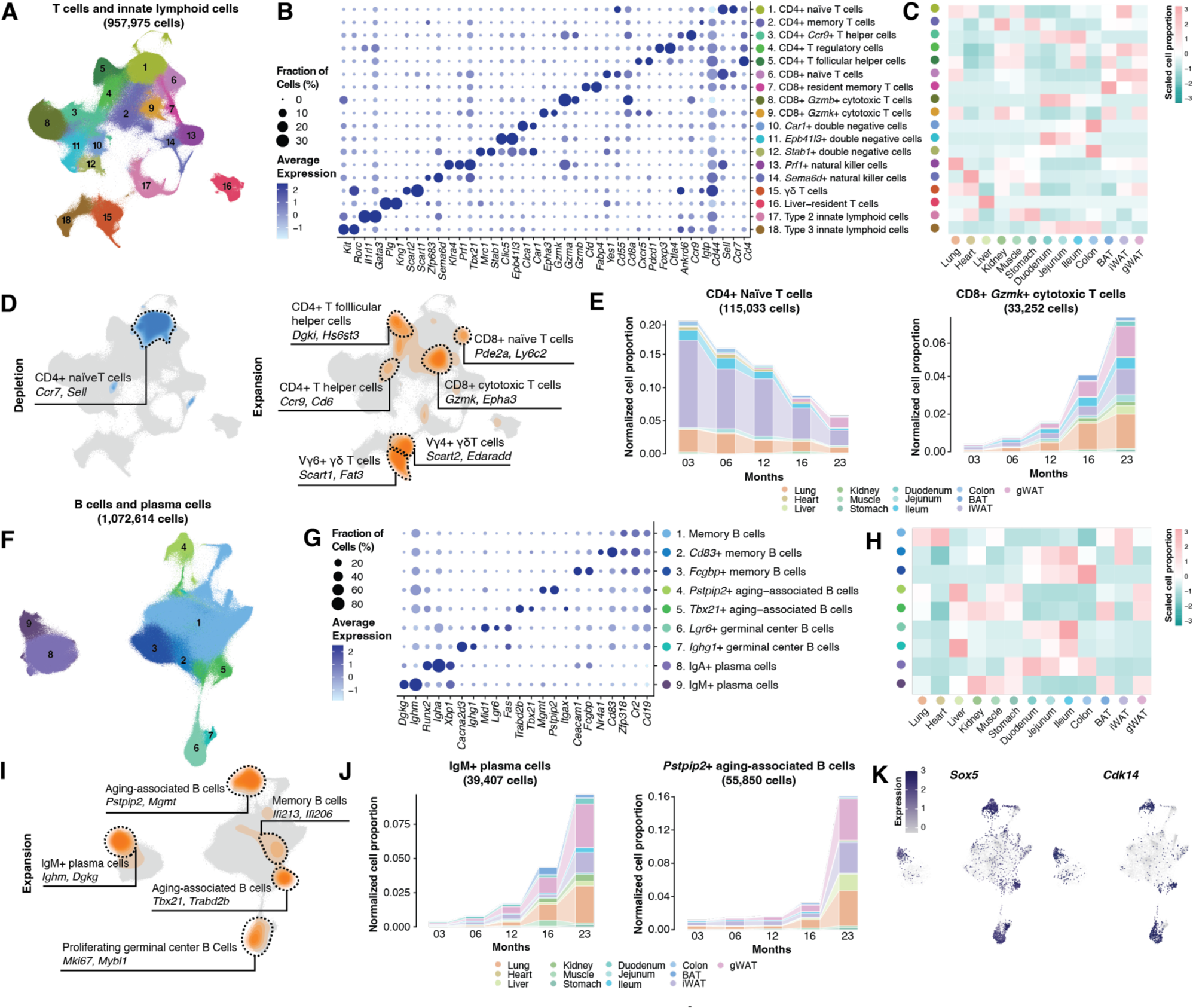
Identifying aging-associated lymphocytes across organs/tissues. (**A**) UMAP visualization of 957,975 T cells and innate lymphoid cells (ILCs) across various organs/tissues, colored by cluster ID. (**B**) Dot plot illustrating marker gene expression for T cell and ILC subtypes. The color denotes average expression values, and the dot size indicates the percentage of cells expressing these markers. (**C**) Heatmap displaying the normalized and scaled distribution of each T cell and ILC subtype across different organs/tissues. (**D**) Density plot highlighting the distribution of significantly depleted (Left) and expanded (Right) T cell and ILC sub-clusters in aging, with their respective marker genes. (**E**) Stacked bar plot depicting the proportion of CD4+ Naïve T cells (Left) and CD8+ *Gzmk+* cytotoxic T cells (Right) within each organ/tissue in wild-type cells, normalized by organ and age group. (**F**) UMAP visualizations of 1,072,614 B cells and plasma cells across organs/tissues, colored according to cluster ID. (**G**) Dot plot showing expression of marker genes for B cell and plasma cell subtypes, with color indicating average expression and dot size reflecting cell expression percentage. (**H**) Heatmap illustrating the normalized and scaled distribution of each B cell and plasma cell subtype across organs/tissues. (**I**) Density plot revealing the distribution of aging-associated B cell and plasma cell sub-clusters with significant expansion in aging, annotated with distinct marker genes. (**J**) Stacked bar plot indicating the expansion of IgM+ plasma cells (Left) and *Petpip2*+ aging-associated B cells (Right) in each wild-type organ/tissue, normalized by organ and age group. (**K**) UMAP visualization demonstrating the widespread expression of *Sox5* and *Cdk14* in expanded B cell and plasma cell populations.

Investigating aging-associated dynamics in T cell subsets, we noted that nearly all T cell clusters that diminished with age could be traced back to the CD4+ naïve T cells (**Fig. 4D**). This trend was consistent across different organs and aligned with previous studies(*38*) (**Fig. 4E, left**). However, age-associated T cell expansion was more varied across distinct molecular states. Expanding cell subtypes included CD4+ T follicular helper cells (*Dgki*+ *Hs6st3*+), CD4+ T helper cells (*Ccr9+* CD6+), CD8+ *Gzmk*+ cytotoxic T cells, CD8*+ Pde2a+ Ly6c2+* T cells, and γδ T cells (**Fig. 4D**). While the CD8*+ Pde2a+ Ly6c2+* T cells were not well characterized in prior studies, its age-related expansion aligns with observations of an age-associated surge in *Ly6c*-expressing immune cells in both bone marrow and spleen (*45*, *46*). This increase may be attributed to a phenomenon wherein proliferating naive CD8+ T cells, in the absence of specific antigen recognition, progressively express *Ly6c*(*47*). Additionally, we observed a broad expansion of CD8+ *Gzmk+* cytotoxic T cells across various organs (**Fig. 4E, right**), in line with prior studies reporting age-related production of pro-inflammatory molecules (*e.g.,* granzyme K) involved in tissue remodeling in aged mice(*44*) and higher abundance of GZMK-expressing CD8*+* T cells in aged human blood(*48*). Interestingly, while the expansion of this CD8*+ Gzmk+* T cell subset occurs at various locations (*e.g.,* kidney, lung, and adipose tissue), its presence is minimal in the intestine, which is dominated by the CD8+ *Gzmb+* T cell subset, suggesting a unique immune-mediated regulatory mechanism in intestinal aging.

Analyzing 1,072,614 pan-organ B cells and plasma cells, we detected nine distinct cell clusters, each marked by unique gene markers and organ-specific distributions (**Fig. 4F-H**). Notably, IgA*+* plasma cells (Cluster 8) were predominantly found in the digestive system (**Fig. 4H**), aligning with their pivotal role in producing immunoglobulin A, a primary defense for the mucosal epithelium against pathogens and toxins(*49*). Similar organ-specific distribution was seen in subsets of memory B cells with high expression of *Cd83* and *Fcgbp* (Cluster 2 and 3) and germinal center B cells (Cluster 6) (**Fig. 4H**). *Cd83* is associated with activated B cells during germinal center reactions(*50*), while IgGFc-binding protein (*Fcgbp*) underpins mucosal immunity in the intestinal lining(*51*). In contrast, other B cells and plasma B cell subtypes were dispersed across multiple organs, such as kidney, lung, and adipose tissue (**Fig. 4H**).

In parallel with age-associated T cell expansion, various B cell subsets were significantly expanded during the aging process (**Fig. 4I**). The first age-associated B cell subset, with increased *Tbx21* expression, resembles a previously reported B cell subset associated with lupus-like autoimmunity in mice(*38*, *52*). The second age-associated B cell subset displays elevated *Pstpip2* expression, a factor linked to macrophage activation, neutrophil migration, and autoinflammatory diseases(*53*) (**Fig. 4J**). Additionally, we detected the age-associated expansion of an IgM+ plasma cell subtype (**Fig. 4J**), marked by elevated *Xbp1, Dgkg,* and *Igbm* expressions. This subtype was widespread in aged mice tissues, including the liver and the adipose tissue (**Fig. 4H**). Unlike other organs, the aged intestine is uniquely featured with a notable rise in *Mki67*+ *Mybl1*+ germinal center B cells. The age-associated proliferation of distinct B and T cell subtypes in the intestine highlights its differential aging process compared to other solid nonlymphoid tissues. We further delved into the molecular programs underlying these expanded B cell populations. Despite varying cellular characteristics and originating organs, they shared the same gene markers like *Sox5* and *Cdk14* (**Fig. 4K**), both involved in the cell cycle and proliferation(*54*, *55*), indicating their roles in age-associated cellular expansion.

### The impact of lymphocyte deficiency on aging

To understand the impact of lymphocytes on cell population dynamics in aging, we employed a “knockdown” approach, targeting lymphocytes throughout the mammalian body using two specific immunodeficient genotypes: *B6.129S7-Rag1tm1Mom/J* and *B6.Cg-Prkdcscid/SzJ*. These models are recognized for lacking functional or mature lymphocytes(*12*, *56*). To validate the lymphocyte deficiency in these models, we compared the main cell populations of the two mutants with age-matched wild-type controls (3-month-old). As anticipated, the majority of the diminished cell populations were lymphocytes, including B cells, T cells, and plasma cells, across various organs (**Fig. 5A**).

**Figure 5.**
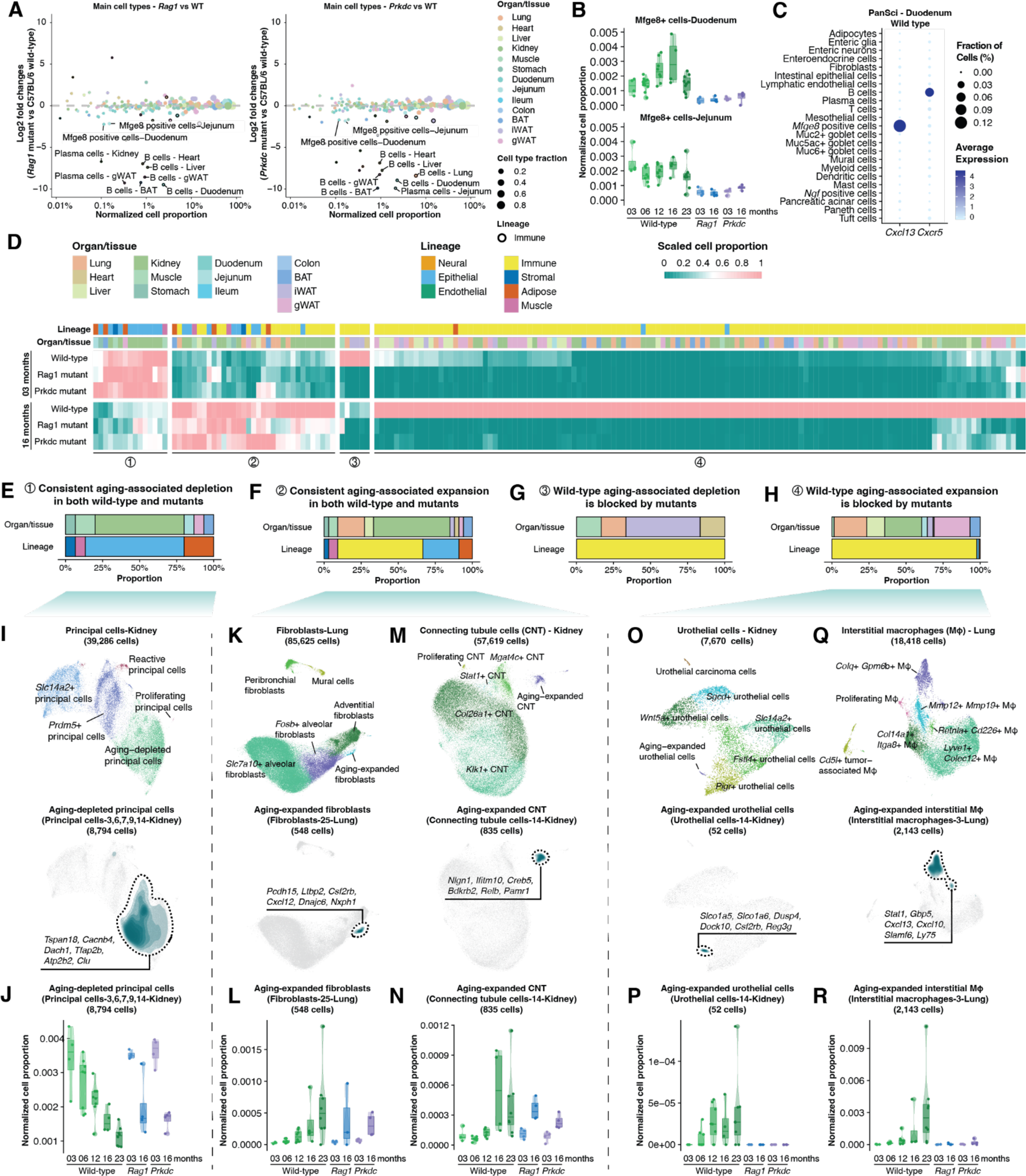
Characterizing lymphocyte-dependent cell population dynamics in aging. (**A**) Scatter plots comparing the proportion changes of main cell types between C57BL/6 wild-type mice and *Rag1* (left) or *Prkdc* (right) mutants. Immune cell lineages are highlighted with black circles, with significant alterations labeled. (**B**) Box plots illustrating the fraction changes of Mfge8+ cells in the duodenum (upper) and jejunum (lower) across life stages in both wild-type and mutant mice. Each dot represents a biological replicate. Box plots display the median (middle line), quartiles (box edges), and 1.5x interquartile range (whiskers). (**C**) Dot plot showcasing the expression of *Cxcl13* and its receptor *Cxcr5* in PanSci’s duodenum dataset, colored by average gene expression and sized by the percentage of cells expressing these markers. (**D**) Heatmap visualizing fraction changes of aging-associated sub-clusters (identified in Figure 3) between 3 and 16 months in C57BL/6 wild-type and immunodeficiency mutants, with hierarchical clustering revealing four distinct dynamic patterns. (**E-H**) Stacked bar plots presenting the proportions of aging-associated sub-clusters from different lineages and organs/tissues in each dynamic pattern. (**I-J**) Case study of kidney principal cells: UMAP visualizations of 39,286 kidney principal cells (I, upper) and density plot depicting the distribution and marker genes of aging-depleted principal cells (J, lower); box plot detailing population shifts in aging-depleted principal cells across different life stages in wild-type and mutant mice (J).(**K-L**) Case study of lung fibroblasts: UMAP visualizations of 85,625 lung fibroblasts (K, upper) and density plot depicting the distribution and marker genes of aging-expanded lung fibroblasts (K, lower); box plot detailing population shifts in aging-expanded lung fibroblasts across different life stages in wild-type and mutant mice (L).(**M-N**) Case study of kidney connecting tubule cells: UMAP visualizations of 57,619 kidney connecting tubule cells (CNT) (M, upper) and density plot showing the distribution and marker genes of aging-expanded CNT (M, lower); box plot detailing population shifts in aging-expanded CNT across different life stages in wild-type and mutant mice (N).(**O-P**) Case study of kidney urothelial cells: UMAP visualizations of 7,670 kidney urothelial cells (O, upper) and density plot showing the distribution and marker genes of aging-expanded urothelial cells (O, lower); box plot detailing population shifts in aging-expanded urothelial cells across different life stages in wild-type and mutant mice (P). (**Q-R**) Case study of lung interstitial macrophages: UMAP visualizations of 18,418 lung interstitial macrophages (Q, upper) and density plot showing the distribution and marker genes of aging-expanded interstitial macrophages (Q, lower); box plot detailing population shifts in aging-expanded interstitial macrophages across different life stages in wild-type and mutant mice (R).

In addition to lymphocytes, we observed a marked decrease in intestine-specific *Mfge8+* follicular dendritic cells (FDCs) (*57*) across two immuno-deficient mutant models. This decline was consistent across anatomical sites (*i.e.,* duodenum and jejunum) and various age stages (**Fig. 5B**). FDCs, recognized for their crucial roles in B cell activation and antibody maturation, are the primary producers of chemokine *Cxc13* in primary follicles and germinal centers of the intestine(*58*). This chemokine, in synergy with the B cell-specific receptor *Cxcr5* (**Fig. 5C**), plays a vital role in B cell positioning within follicles and is essential in defining the secondary lymphoid tissue architecture, including lymph nodes and Peyer’s patches(*59*). Noteworthily, our findings reveal that FDC-lymphocyte interactions are crucial for sustaining this intestinal stromal cell population.

Extending down to the sub-cluster level, we identified 289 sub-clusters exhibiting significant population changes in two immuno-deficient models (**fig. S12)**. As expected, the depleted sub-clusters are primarily associated with lymphocytes across various organs. Interestingly, several sub-clusters significantly increased upon lymphocyte-knockdown (*e.g., Rnf213+ Ddx60+* intestinal epithelial cells in duodenum and jejunum), suggesting that lymphocytes might play a role in limiting the growth of these intestinal epithelial cells. Meanwhile, specific stromal sub-clusters (*e.g., Serpine1+ Jun+* adipocytes in the lung) were depleted in both immuno-deficient mutant models, hinting at potential stromal-immune crosstalks that warrant future exploration.

We next clustered aging-associated cell subtypes based on their dynamics in two immunodeficient models (**Fig. 5D**), focusing on subtypes that are detected and displayed consistent alterations in both mutants. Intriguingly, 15 subtypes with age-associated depletion and 33 subtypes with age-associated expansion exhibited consistent patterns in both wild-type and mutant models, suggesting their population changes are not directly caused by lymphocyte involvement (**Fig. 5, E and F**). Representative examples include the aging-associated depletion of *Tspan18+* kidney principal cells (**Fig. 5I-J**), expansion of *Pcdh15*+ lung fibroblast (**Fig. 5K-L**) and *Nlgn1+* kidney connecting tubule (CNT) cells (**Fig. 5, M and N**). Molecular analysis of these cells offers insights into aging-linked organ dysfunction. For example, the aging-depleted renal principal cells correspond to a group of progenitor cells (*e.g., Dach1, Tfap2b, Tspan18*(*60*)) (**Fig. 5I**) and show elevated expression of genes crucial for calcium homeostasis (*e.g., Cacnb4, Atp2b2*(*61*)). This may indicate a susceptibility to calcium-induced cellular stress, potentially predisposing them to age-related damage and depletion. These findings highlight the complex interplay of cellular changes in aging and suggest mechanisms beyond direct lymphocyte interactions.

In contrast, other aging-associated subtypes presented distinct population dynamics between wild-type and mutant models, indicative of immune-dependent regulation (**Fig. 5D**). A considerable proportion (128 out of 138) of these subtypes were lymphocytes, impacted by their depletion in the mutants (**Fig. 5, G and H**). For instance, the aging-associated depletion of naive T cells and expansion of most lymphocytes were absent in the aged mutant. In addition, we observed several non-lymphocyte cell populations that displayed altered dynamics in both mutants. A notable rescued cell-type-specific expansion is a *Slco1a5*+ kidney urothelial subtype (*i.e.,* urothelial cells-14) featured with an enriched expression of genes indicative of immune stimulation (*e.g., Csf2rb*(*62*), **Fig. 5, O and P**). Likewise, the expansion of a unique subtype of *Colq*+ lung interstitial macrophage (*i.e.,* interstitial macrophage-3) was halted in both mutants (**Fig. 5, Q and R**). This macrophage subtype is characterized by genes associated with lymphocytes interaction (*e.g., Cxcl13*, *Cxcl10*), illuminating the critical role of lymphocytes in driving its expansion during aging. These observations underscore the instrumental role of lymphocytes in regulating the dynamics of these cell populations. Consequently, targeted ablation of lymphocytes could be a viable strategy for the in-depth functional analysis of cellular interactions throughout the organism.

## Discussion

In this study, we’ve generated an extensive catalog highlighting the intricate dynamics of cell population changes upon aging. This was achieved through high-throughput single-cell transcriptome analysis of over 20 million cells from 623 tissue samples spanning various life stages (3, 6, 12, 16, 23 months), sexes, and genotypes. The analysis revealed a complex and dynamic landscape of aging at the cellular level, uncovering more than 10 main cell types and over 200 subtypes undergoing significant age-associated depletion or expansion. Notably, while some cell types, such as lymphocytes, have previously been documented to expand with age, our study has uncovered a range of rare cellular states that remain underexplored, such as the depletion of *Tspan18*+ principal cells and the expansion of *Nlgn1*+ connecting tubule cells within renal tissues. These findings were consistently observed across varying ages and even genotypes, underscoring their potential as anti-aging targets for further therapeutic exploration. Additionally, we discovered 73 subclusters that are highly sex-specific across different ages, as well as sexually dimorphic cellular dynamics in aging, exemplified by the accelerated decline of *Mirg*+ muscle cells in females.

Moreover, our data suggest that aging at the cellular level unfolds through a series of dynamic waves rather than following a simple linear trajectory akin to the patterns observed with the DNA Methylation Clock(*63*). Early stages (3 to 12 months, mirroring human ages 20 to 42) are primarily characterized by the depletion of specific cell types within the adipose, muscle, and epithelial lineages. In contrast, later stages (12 to 23 months, analogous to human ages 42 to 68) are dominated by a pronounced expansion of various immune cell populations. This observation challenges the traditional Wear-and-Tear Theory of aging(*64*), proposing instead that a complex array of regulatory mechanisms are at play. These mechanisms orchestrate a series of coordinated cell population transitions, which unfold throughout the aging process and vary distinctly between each examined age interval. Our findings also align with prior reports(*65*), highlighting the advantages of initiating anti-aging interventions in early life, given cellular depletion occurs in the initial stages of aging.

Furthermore, our study has uncovered organ-specific changes within broadly distributed cell types (*e.g.,* immune cells) through a comprehensive analysis of cell population shifts across various organs. For example, we observed a consistent decrease in CD4+ naïve T cells and an increase in CD8+ *Gzmk+* cytotoxic T cells and age-related B cell subsets. These consistent patterns suggest a universal regulatory mechanism governing immune aging throughout the body. Notably, specific organ systems displayed distinct aging dynamics, with early immune expansion occurring predominantly in the kidney and lung, while later expansions were observed in the liver and others, potentially linked to the onset of organ-specific aging-associated diseases. The intestinal environment, in particular, displayed a unique profile, with an increase in specific aging-associated B cells (*e.g., Mki67+ Mybl1+* germinal center B cells) and T cell subtypes (*e.g.,* CD8+ *Gzmb+* T cells), highlighting a potentially unique aspect of immune regulation in gut aging relative to other nonlymphoid tissues.

Utilizing a “cell-knockdown” strategy analogous to “gene-knockdown” in functional genomics, we targeted lymphocytes to interrogate their role in the aging-related population dynamics of other cell types. This strategy was instrumental in delineating the complex interplay between lymphocytes and other cell types, evidenced by the halted increase of *Slco1a5*+ kidney urothelial cells and *Colq*+ lung interstitial macrophages upon lymphocyte reduction. However, efforts to restore most depleted cell populations through immune knockdown proved largely unsuccessful, reinforcing the temporal hierarchy of cellular depletion preceding expansion. This underscores the intricate and multi-layered regulatory mechanisms governing cell population changes throughout aging.

Of note, the scalability of the single-cell combinatorial indexing strategy has been pivotal to our study’s success, as it allows the inclusion of multiple individuals, with a sex balance, at various aging stages, with all cells from each organ profiled concurrently. This is a significant advancement over traditional approaches that often require the integration of different technical batches. While our primary focus has been on cell population dynamics throughout aging, the applicability of our dataset opens avenues for investigating a myriad of compelling biological questions, from cell-type-specific transcriptomic alterations associated with aging to variations in cellular profiles due to differences in sexes and genotypes. Furthermore, the depth of our dataset, featuring single-nucleus gene expression with full gene body coverage, allows for exploring cell type-specific dynamics concerning isoform variation or non-coding RNA expression changes during aging.

In summary, our work has meticulously charted an extensive spectrum of over 3,000 unique cellular states in the mammalian system, identifying over 200 that exhibit significant aging-related changes in a tightly coordinated manner. We uncovered lymphocyte-dependent cellular population shifts associated with aging by harnessing scalable single-cell genomic techniques alongside mutant strain analysis. This ‘’Cell-omics’ strategy—mirroring the progress made in high-throughput genomic sequencing—sets the stage for identifying key cellular targets and their regulatory network in various aging-related conditions, which holds the potential to spur therapeutic innovations aimed at restoring cellular functions and rejuvenating the systemic biological processes of organisms in aging and diseases.

## Acknowledgments

We thank members of the Cao lab for helpful discussions and feedback. We also thank members of the Rockefeller University High Performance Computing Core, Comparative Bioscience Center, and Flow Cytometry Resource Center for their invaluable support.

## Funding

This work was funded by grants from NIH (DP2HG012522, R01AG076932, and RM1HG011014 to J.C.) and the Sagol Network GerOmic Award for J.C. The project was partly supported by a Longevity Impetus Grant from Norn Group. B.W. and M.H. were funded by NIH/NIMH RF1MH132662, CIRM DISC0-14514 and NIH/NHGRI U24HG002371. R.S. was funded by RM1HG011014-02 and 1OT2OD033760-01.

## Author contributions

J.C. and W.Z. conceptualized and supervised the project; Z.Z. optimized nuclei extraction methods, with input from J.L.; Z.Z. performed mouse dissection, nuclei extraction, single-cell RNA-seq experiment, with assistance from C.S. and W.J.; Z.Z. performed computational analyses with insights from Z.Lu, A.S, and A.A.; B.W. and M.H. built the UCSC cell browser for PanSci; Z.Li, G.M. and R.S. built the Azimuth application for PanSci; J.C., W.Z. and Z.Z. wrote the manuscript with input from all co-authors.

## Competing interests

In the past 3 years, R.S. has received compensation from Bristol-Myers Squibb, ImmunAI, Resolve Biosciences, Nanostring, 10x Genomics, Neptune Bio, and the NYC Pandemic Response Lab. R.S. is a co-founder and equity holder of Neptune Bio.

## Data and materials availability

A detailed protocol of *EasySci* is outlined in (*14*). The computational processing pipeline for read alignment and gene count matrix generation is available on GitHub at this repository: https://github.com/JunyueCaoLab/EasySci. Raw FASTQ files, processed count matrices, cell metadata, and gene metadata can be downloaded from NCBI GEO under accession number GSE247719. PanSci data can be interactively accessed in UCSC cell browser at https://mouse-pansci.cells.ucsc.edu and mapped in Azimuth application at https://app.azimuth.hubmapconsortium.org/app/mouse-pansci.

## Supplementary Materials

### Materials and Methods

#### Animals and organ collection

C57BL/6 wild-type mice were obtained from the Jackson Laboratory and the National Institution on Aging colony at Charles River. The immunodeficient strains, *B6.129S7-Rag1tm1Mom/J* (JAX #002216) and *B6.Cg-Prkdcscid/SzJ* (JAX #001913), were also obtained from the Jackson Laboratory. All strains were housed according to standard protocols, with same sex- and age-matched groups. The sex-balanced cohorts ranged in age from 106 days to 704 days. Comprehensive metadata for each animal, including mouse individual ID, sex, age, birth and euthanasia dates, and body and organ/tissue weights, are detailed in **table S1A**.

All animal procedures were in accordance with institutional, state, and government regulations and approved under the IACUC protocol 21049. In brief, animals of the same age and sex were euthanized, and the organ/tissue collection for each batch was carried out by the same person on the same day, with approximately one-hour intervals between each euthanasia to ensure temporal consistency. This scheduling was designed to reduce the potential circadian rhythm’s effect on transcriptomic data across different sexes and age groups. For each mouse, whole organs/tissues were then dissected in the following order: inguinal adipose tissue (with inguinal lymph nodes), stomach, small intestine (duodenum, jejunum, ileum), colon, perigonadal adipose tissue, kidney, liver, heart, lung, hindlimb muscle, brown adipose tissue. A complete organ/tissue set was profiled for most mouse individuals, with additional specimens included to compensate for any losses during dissection or dissociation processes. For the immunodeficient strains, the duodenum was exclusively profiled to represent the small intestine. All collected organs/tissues are washed thoroughly in ice-cold HBSS (Thermo Fisher #14175095) and immediately flash-freeze in liquid nitrogen. Snap-frozen tissues are manually pulverized on dry ice with a chilled hammer, aliquoted, and stored in liquid nitrogen until further processing.

#### Nuclei extraction from multiple mammalian organs

The 10X PBS-hypotonic stock solution was prepared using the method described in reference (*15*). On the day of nuclei extraction, 1X hypotonic lysis buffer was freshly prepared by diluting the 10x stock solution with RNase-free water (Corning, #46-000-CM), supplemented with 3mM MgCl2, 1% Diethyl pyrocarbonate (Sigma Aldrich, #40718) and specifically optimized detergent for each organ/tissue type: 0.025% IGEPAL CA-630 (VWR, #IC0219859650) for kidney, lung, liver, brown adipose tissue, inguinal adipose tissue, and perigonadal adipose tissue; 0.01% Digitonin (Thermo Fisher, #BN2006) for heart, muscle, duodenum, jejunum, ileum, and colon; 0.49% CHAPS (Sigma Aldrich, #220201) for the stomach. Additionally, 0.33M sucrose (Sigma Aldrich, #S0389) was included in the working lysis buffer for the stomach and intestinal tissues (duodenum, jejunum, ileum, and colon) to maintain osmotic balance and protect the nuclei.

For nuclei extraction, dry tissue powder stored in liquid nitrogen was quickly transferred into 10 mL of a pre-prepared lysis solution. After a brief 10-second vortex to disperse large chunks, the mixture underwent a 15-minute incubation at 4°C with constant rotating. It was then strained through a 40 μm cell strainer (VWR, #470236-276) using a 5 mL syringe plunger, with an additional 5 mL of lysis solution used to rinse the filter. The extracted nuclei were collected by centrifugation at 500g for 5 minutes at 4 °C and resuspended in a nuclei suspension buffer. This buffer contained 10 mM Tris-HCl pH 7.5 (Thermo Fisher, #15567027), 10 mM NaCl (Thermo Fisher, #AM9760G), 3 mM MgCl2 (Sigma Aldrich, #68475-100ML-F), 1% SUPERase⋅In RNase Inhibitor (Thermo Fisher, #AM2696), and 0.2 mg/mL BSA or Recombinant Albumin (New England Biolabs, #B9000S or #B9200S), supplemented with 0.005 mg/mL DAPI (Thermo Fisher, #D1306) for fluorescence-activated cell sorting (FACS). FACS was performed on a SH800 Cell Sorter with a 100 μm sorting chip (Sony, #LE-C3210), aiming to include all DAPI-positive singlet nuclei, which aids in recovering the global cell population while removing cellular debris and doublets. Nuclei were collected into a 1.5 mL tube (Eppendorf, #022431021) containing 100 μL of nuclei suspension buffer and subsequently concentrated by centrifugation at 500g for 5 minutes at 4 °C.

We introduced a control step to assess batch effects during library preparation and sequencing. Specifically, control kidney nuclei, extracted from pooled mouse kidney samples using the previously described methods, were spiked into each library at the reverse transcription stage. These control nuclei were aliquoted into 1.5 mL tubes and underwent a slow freeze in a nuclei suspension buffer with an added 10% DMSO (VWR, #97063-136), stored at −80 °C. When required for sorting, an aliquot of these control nuclei was rapidly thawed in a 37 °C water bath and then sorted in conjunction with the actual experimental samples.

#### *EasySci* library construction and sequencing

The sequencing library generation for the sorted nuclei was conducted in accordance with the *EasySci* protocol (*14*). Initially, the sorted nuclei were allocated to 96-well plates (Geneseesc, #24-302) for reverse transcription. Here, both indexed oligo-dT and indexed random hexamer primers were employed to introduce the first index. Subsequently, these nuclei underwent pooling, washing, and re-distribution into new 96-well plates for the second index attachment via ligation. This was followed by another set of pooling and washing, after which the nuclei were placed into final plates for second-strand synthesis and purification. The concluding steps were tagmentation with Tn5 transposase and PCR for the final index addition. The final PCR products were then pooled and purified using a 0.8X volume of AMPure XP SPRI Reagent (Beckman Coulter, #A63882). Library quality was verified using an Agilent TapeStation, and sequencing was performed on an Illumina NovaSeq 6000 System with twenty-five S4 flow cells. Read alignment and gene/exon count matrix generation for the single-cell RNA-seq were performed using the pipeline we developed for *EasySci* (*14*). Control kidney samples from each run, identified by reverse transcription barcodes, were compiled to create a gene count matrix, enabling the assessment of batch effects.

#### Cell filtering, clustering, and marker gene identification

Cells from each NovaSeq run are split into an organ/tissue-specific count matrix for data cleaning. Briefly, the low-quality cells, merged from oligo-dT reads and random hexamer reads, were filtered out if they met one of the following criteria: 1) unmatched rate (proportion of reads not mapping to any exon or intron) >= 0.4, 2) UMI count < 200, 3) gene count < 100. Then, Scrublet (version 0.2.3) (*66*) was applied to each count matrix with parameters (min_count = 3, min_cells = 3, vscore_percentile = 85, n_pc = 30, expected_doublet_rate = 0.08, sim_doublet_ratio = 2, n_neighbors = 30. Cells with doublet scores over 0.2 were annotated as doublets and discarded.

The count matrices from each NovaSeq run were aggregated to form organ/tissue-specific matrices. For each organ/tissue, main cell type clustering was carried out using Scanpy (version 1.9.3) (*67*) through the following steps: 1) Normalization of the gene count matrix per cell by total UMI count followed by logarithmic transformation. 2) Selection of the 5000 most variable genes per organ/tissue matrix, scaling their expression to zero mean and unit variance. 3) Dimension reduction using PCA, utilizing the top 50 principal components to construct a neighborhood graph (n_neighbor = 50). 4) Leiden clustering (resolution = 0.5). 5) Further dimension reduction with UMAP into 2D space (min.dist = 0.01). Differentially expressed genes within clusters for each organ/tissue were identified using the differentialGeneTest() function in Monocle2 (version 2.28.0) (*68*). Specific gene markers were selected based on differential expression across clusters, with criteria including a maximum 5% false discovery rate, a minimum 2-fold expression difference between top-ranked and second-top clusters, and TPM over 50 in the highest-ranked cluster. Clustering was refined by merging adjacent clusters if they had few differentially expressed genes or shared high expression of the same literature-nominated marker genes. Main cell type annotations were based on organ/tissue-specific published cell type markers. This strategy enabled the recovery of almost all main cell types identified in similar atlasing studies, accommodating variations in species, developmental stages, and methodologies. We also identified 16 unknown cell types in certain tissues, labeling them according to their top enriched differentially expressed gene markers specific to each tissue.

For cross-organ cell clustering from wild-type samples, we combined wild-type cells from our dataset with the previously generated brain dataset (*14*). The Scanpy pipeline was reapplied for dimension reduction and clustering (n_top_genes=2000, n_neighbors=50, n_pcs=50, min_dist=0.15, resolution=0.5), and lineages were manually annotated in 2D UMAP space.

#### Sub-clustering analysis

To identify sub-clusters within each main cell type with higher resolution, we employed a similar pipeline described in the previous study (*14*). Briefly, each gene count matrix and exon count matrix for each main cell type is normalized, log-transformed, and scaled. These matrices were then subjected to PCA, from which the top 30 principal components of the gene count matrix and the top 10 from the exon count matrix were extracted and combined into a single matrix. This combined matrix underwent further processing through Leiden clustering and dimension reduction using UMAP. To identify the enriched genes for each sub-cluster, we computed the aggregated gene expression per sub-cluster and prioritized the prominently expressed sub-clusters for each gene. A gene specificity score was calculated to assess the uniqueness of gene expression in the most expressed sub-cluster. This dual filtering approach enables a swift and comprehensive assessment of the genetic landscape of each sub-cluster. Enriched genes for sub-clusters of interest were further validated by differentially expressed gene analysis through Monocle2 (*68*).

For case studies illustrated in Figures 3, 4 and 5, UMAP coordinates were calculated based on gene count matrix alone. In Figure 3, all wild-type cells from identified 230 aging-associated sub-clusters were extracted; in Figure 4, all cells annotated as immune lineage were selected, re-clustered, and classified into 1) T cells and innate lymphoid cells, 2) B cells and plasma cells, and 3) myeloid cells for further annotation; in Figure 5, main cell types to which the targeted sub-cluster belonged was isolated. Cells selected for each figure were then subjected to the above-mentioned clustering pipeline. Differentially expressed genes for each cluster within the selected cell group were identified with the differentialGeneTest() function in Monocle2 (version 2.28.0) (*68*), applying a filter criteria that includes 1) a maximal false discovery rate of 5%, 2) a minimum 2-fold expression difference between the top two ranked sub-cluster, and 3) TPM greater than 50 in the highest-ranked sub-cluster. Clusters were merged if they shared the same marker genes. Distinct clusters expressing marker genes of other cell types are further excluded as potential doublets.

#### Intra-dataset cross-validation analysis

To confirm the accuracy of the main cell types and sub-cluster annotation, we implemented a general-purpose support vector machine classifier for intra-dataset cross-validation, mirroring the methods outlined in the reference (*69*). Briefly, we randomly sampled up to 2,000 cells from each cell type, or all cells for cell types with fewer than 2,000 cells. For main cell type validation, we combined sampled cells from the same organ or tissue; for sub-cluster validation, we combined sampled cells from each main cell type. These were then used as input for a 5-fold cross-validation using an SVM classifier with a linear kernel. The complete gene count transcriptome was utilized for predicting both main cell types and sub-clusters. The specificity of our cell type annotation was assessed by calculating the cross-validation F1 score. As a control, we randomly permuted the cell type labels and subjected them to the same analysis pipeline.

#### Identifying aging-associated dynamic waves

To assess cell population dynamics across different conditions, including age groups, sex, and genotype, at both the main cell type and sub-cluster levels, we generated organ/tissue-specific cell count matrices across mouse individuals (cell type X individual). Cell numbers of each cell type for individual mice (serving as replicates) in a particular organ or tissue were counted and then normalized against the total cell number obtained from the corresponding organ or tissue of each individual mouse. Likelihood-ratio test was employed for identifying differentially abundant cell types using the differentialGeneTest() function of Monocle2 (version 2.28.0) (*68*). For fold change calculations, we first normalized the number of cells in each cell type relative to the total cell count in the respective condition. We then compared these normalized values between the case and control conditions, incorporating a small numerical value (10^−6^) to reduce the noise from very small clusters.

To classify a main cell type or sub-cluster as a “significantly changed cell type,” we set specific criteria: 1) a maximum false discovery rate of 0.05 and 2) a fold change higher than 2 between conditions. Additionally, we established more stringent criteria for identifying aging-associated cell types that show consistent changes across the aging process. We focused on two age intervals — “16 months vs 3 months” and “23 months vs 6 months” — and performed differential abundance tests separately for each interval. A main cell type or sub-cluster was considered an “aging-associated cell type” if it met the following conditions: 1) significant changes in both intervals (q-value 16v3 < 0.05, q-value 23v6 < 0.05), 2) a fold change at least 2 in both intervals (absolute(fold-change 16v3) ≥ 2, absolute(fold-change 23v6) ≥ 2), and 3) consistent dynamic directions between two age intervals.

To identify the aging-associated dynamic waves, we generated a cell count matrix across five life stages (cell type X time points). Cell numbers of each cell type for each time point were counted and then normalized against the total cell number from the corresponding organ or tissue of each time point. The cell count matrix across ages for identified “aging-associated cell type” was extracted and subject to hierarchical clustering. Each cluster was manually inspected and categorized into each aging-associated dynamic wave.

#### Identifying sex-specific and genotype-specific cell types

Similar to identifying aging-associated cell types, we constructed organ/tissue-specific cell count matrices across mouse individuals (cell type X individual) and applied a likelihood-ratio test through differentialGeneTest() function in Monocle2 (version 2.28.0) (*68*) under specific conditions. For identifying sex-specific cell types, we compared the differential cell abundance between female and male individuals within each age group independently. Cell types were designated as “sex-specific” based on the following criteria: 1) a maximum false discovery rate of 0.05; 2) a minimum fold change of 2 between sexes; 3) sex-specificity is consistent across five age groups. For identifying genotype-specific cell types, two lymphocyte-deficient mutant strains were treated as biological replicates. Our analysis focused on cell types demonstrating consistent alterations between two mutants. We first compare the differential cell abundance between each mutant and wildtype at 3 months and 16 months. Cell types were designated as “genotype-specific” based on the following criteria: 1) a maximum false discovery rate of 0.05; 2) a minimum fold change of 2 between the mutant and wildtype; 3) mutant-specific change is consistent in both genotypes and across two assessed age groups.

## Figs. S1 to S13

**Fig. S1.**
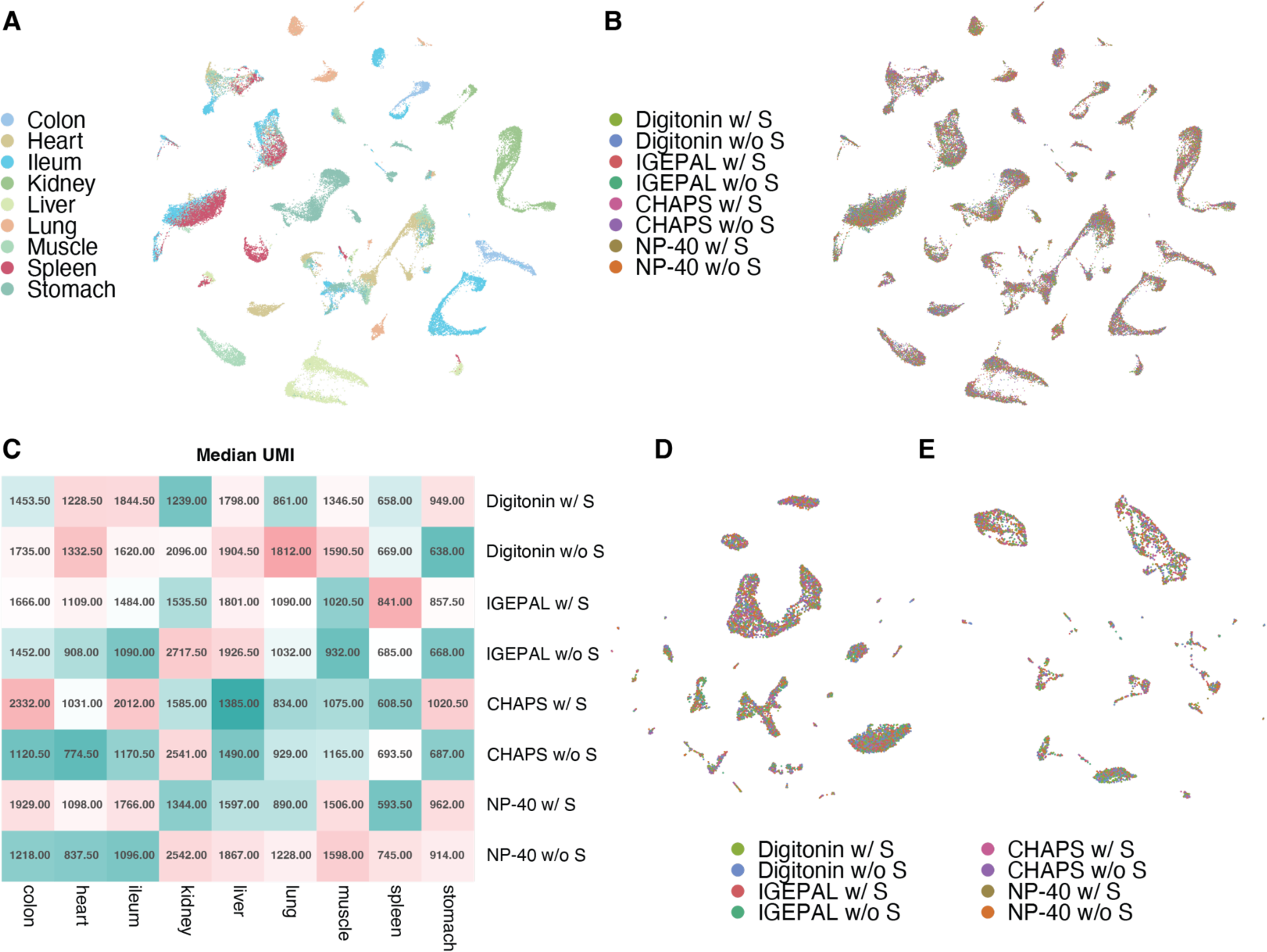
Optimization of lysis conditions for single-cell profiling of diverse mammalian tissues. This figure presents an assessment of lysis conditions tailored for single-cell transcriptome library preparation across a variety of mammalian tissues. (**A-B**) UMAP plots illustrating the clustering results of 49,264 cells, delineated by tissue origin (A) and by lysis conditions (B). It is noteworthy that cells processed in the same hypotonic lysis conditions with different additives demonstrate minimal batch effects without computational integration. Hypotonic lysis buffer working solution is prepared fresh, with specific additives introduced just before nuclei extraction: digitonin, 0.01% digitonin; IGEPAL, 0.025% IGEPAL; CHAPS, 0.49% CHAPS; NP-40, 0.2% NP-40; S, 0.33M sucrose. (**C**) A heatmap representation detailing the median UMI counts retrieved per cell across each tissue type under variable lysis protocols. (**D-E**) Representative UMAP of 10,079 cells from ileum (D), and 4,170 cells from colon (E), colored by lysis conditions. Importantly, the dimensionality reduction analyses were conducted without batch correction for lysis conditions, reinforcing the absence of lysis-condition-induced bias in cell-type representation.

**Fig. S2.**
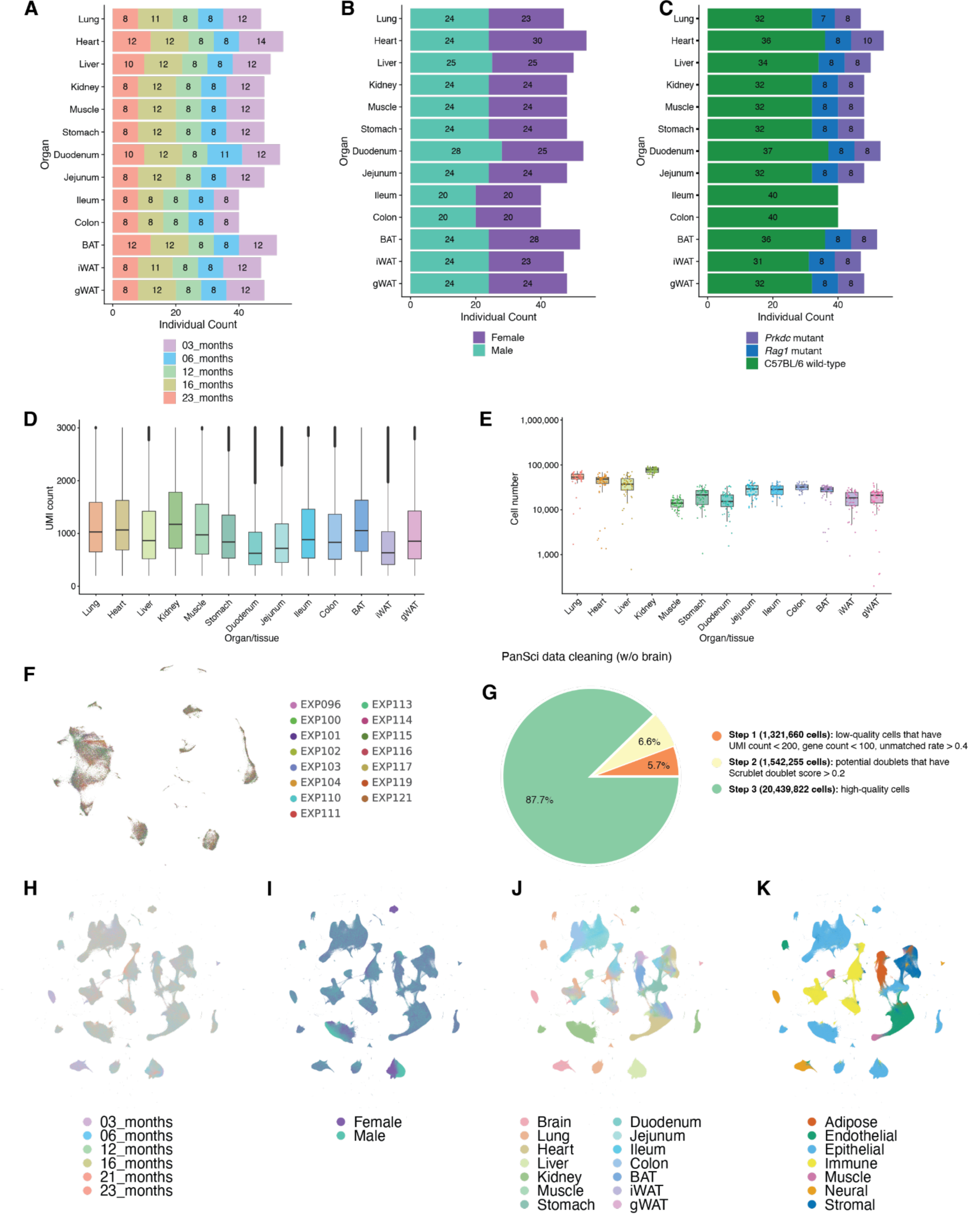
Quality control metrics for the PanSci dataset. (**A-C**) Bar plots showing the number of mouse individuals per organ, colored by age group (A), sex (B) and genotype (C). (**D-E**) Box plot showing the UMI per cell (D) and cell numbers per individual (E) for each organ/tissue of the PanSci dataset without brain. (**F**) UMAP visualization of 112,002 kidney cells spiked in each sequencing library, no data integration was applied. (**G**) Pie chart showing the cell numbers after each of the data cleaning steps. (**H-K**) UMAP visualizations of 15,589,090 wild-type cells colored by age group (H), sex (I), organ/tissue (J) and lineage (K) same as in Figure 1C.

**Fig. S3.**
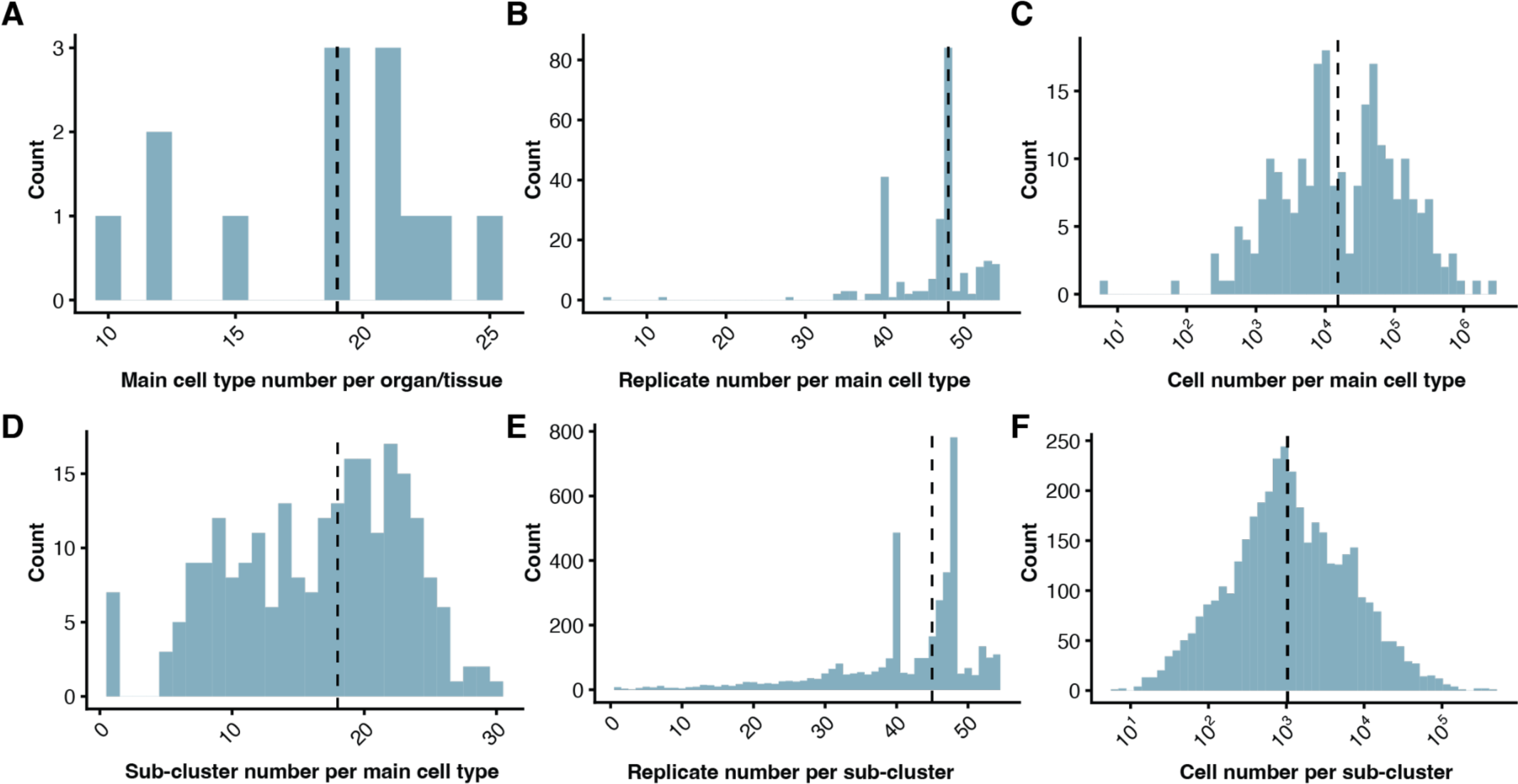
Quality control metrics for the identified main cell types and sub-clusters. (**A-C**) Histogram showing the number of main cell types identified for each organ/tissue (A; median: 19 main cell types per organ/tissue), mouse individual replicates number for each main cell type (B; median: 48 replicates per main cell type), and cell number for each main cell type (C; median: 15,321 cells per main cell type) with a dashed line showing the median number. (**D-F**) Histogram showing the distribution of sub-clusters identified in each main cell type (D; median: 18 sub-clusters per main cell type), mouse individual replicates number for each sub-cluster (E; median: 45 replicates per sub-cluster), and cell number for each sub-cluster (F; median: 1,035 cells per sub-cluster) with a dashed line showing the median number.

**Fig. S4.**
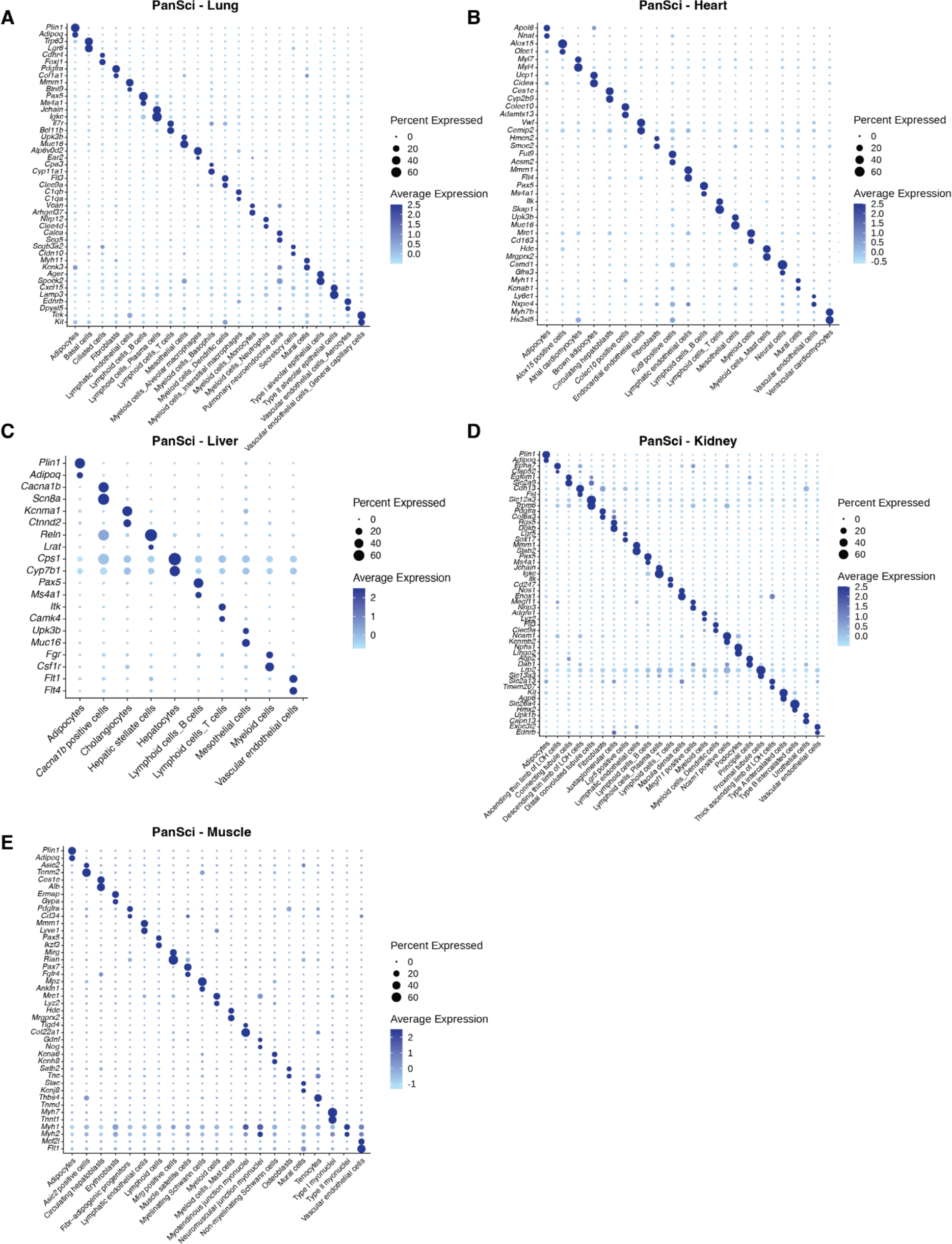
Characterization of main cell types in lung, heart, liver, kidney, and muscle. (**A-E**) Dot plot illustrating gene marker expression for annotating main cell types in lung (A), heart (B), liver (C), kidney (D), and muscle (E) for PanSci. The color denotes average expression values, and the dot size indicates the percentage of cells expressing these markers.

**Fig. S5.**
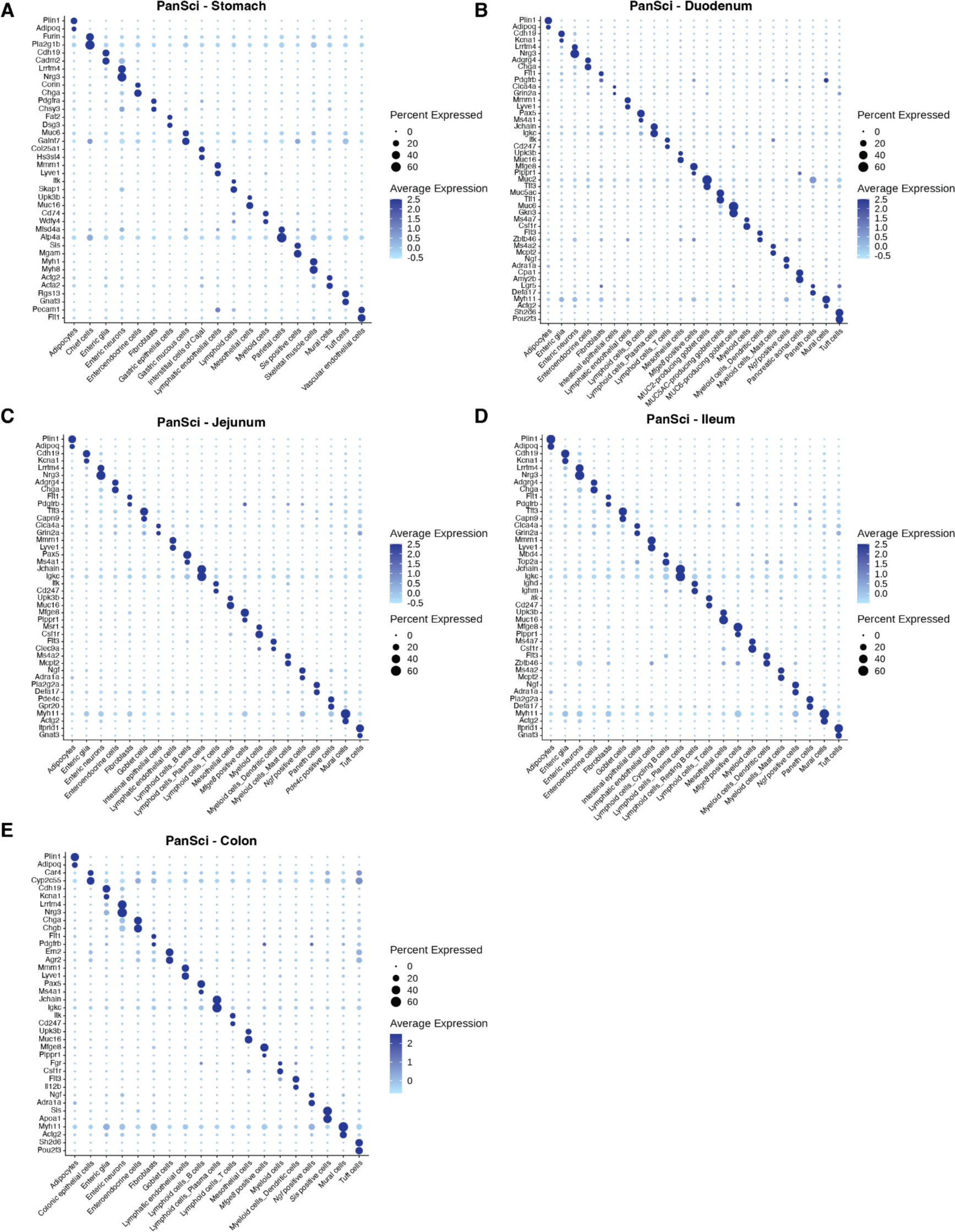
Characterization of main cell types in organs/tissues of the gastrointestinal tract. (**A-E**) Dot plot illustrating gene markers’ expression for annotating main cell types in stomach (A), duodenum (B), jejunum (C), ileum (D), and colon (E) for PanSci. The color denotes average expression values, and the dot size indicates the percentage of cells expressing these markers.

**Fig. S6.**
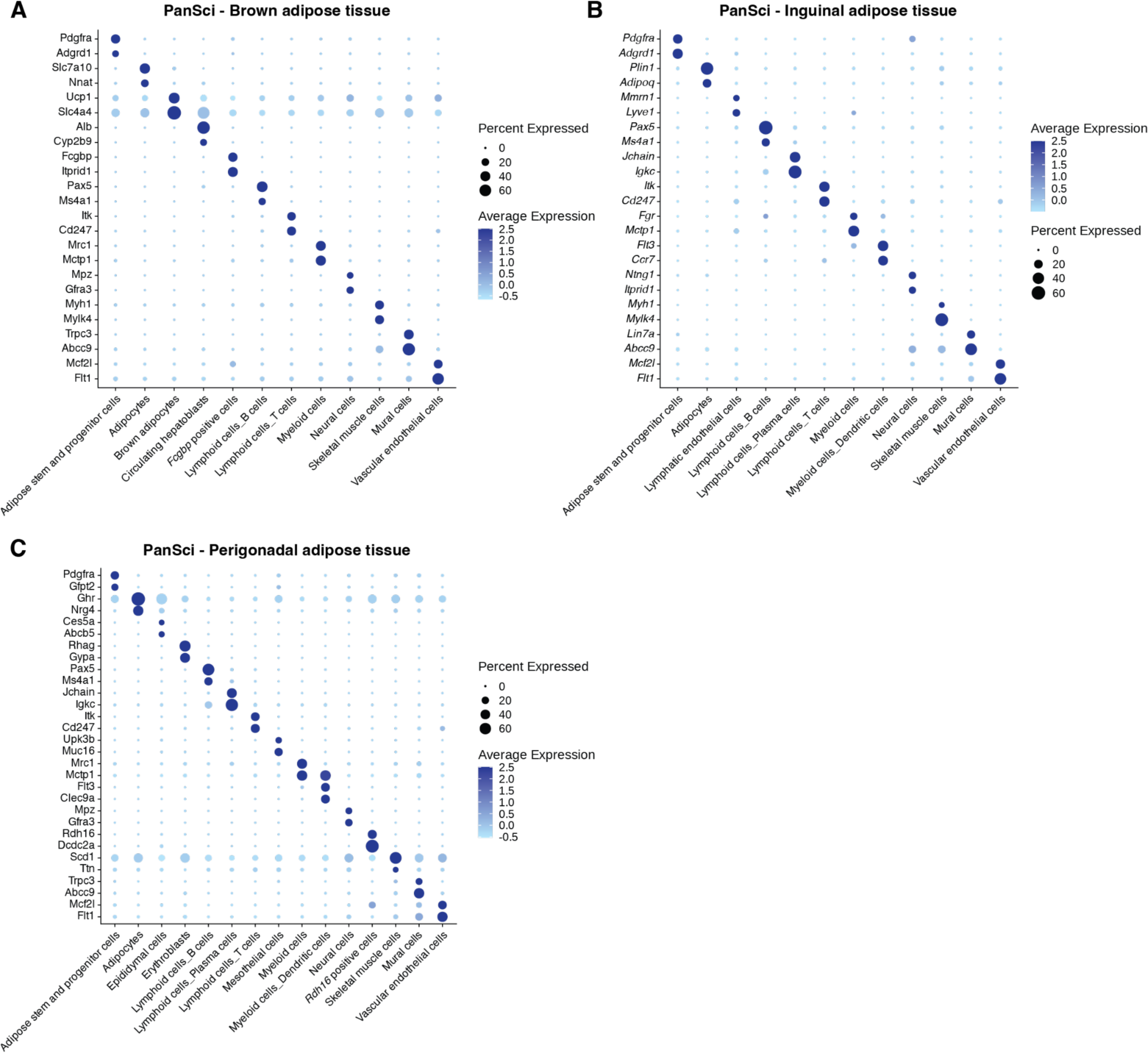
Characterization of main cell types in adipose tissues. (**A-C**) Dot plot illustrating gene markers’ expression for annotating main cell types in brown adipose tissue (A), inguinal adipose tissue (B), and perigonadal adipose tissue (C) for PanSci. The color denotes average expression values, and the dot size indicates the percentage of cells expressing these markers.

**Fig. S7.**
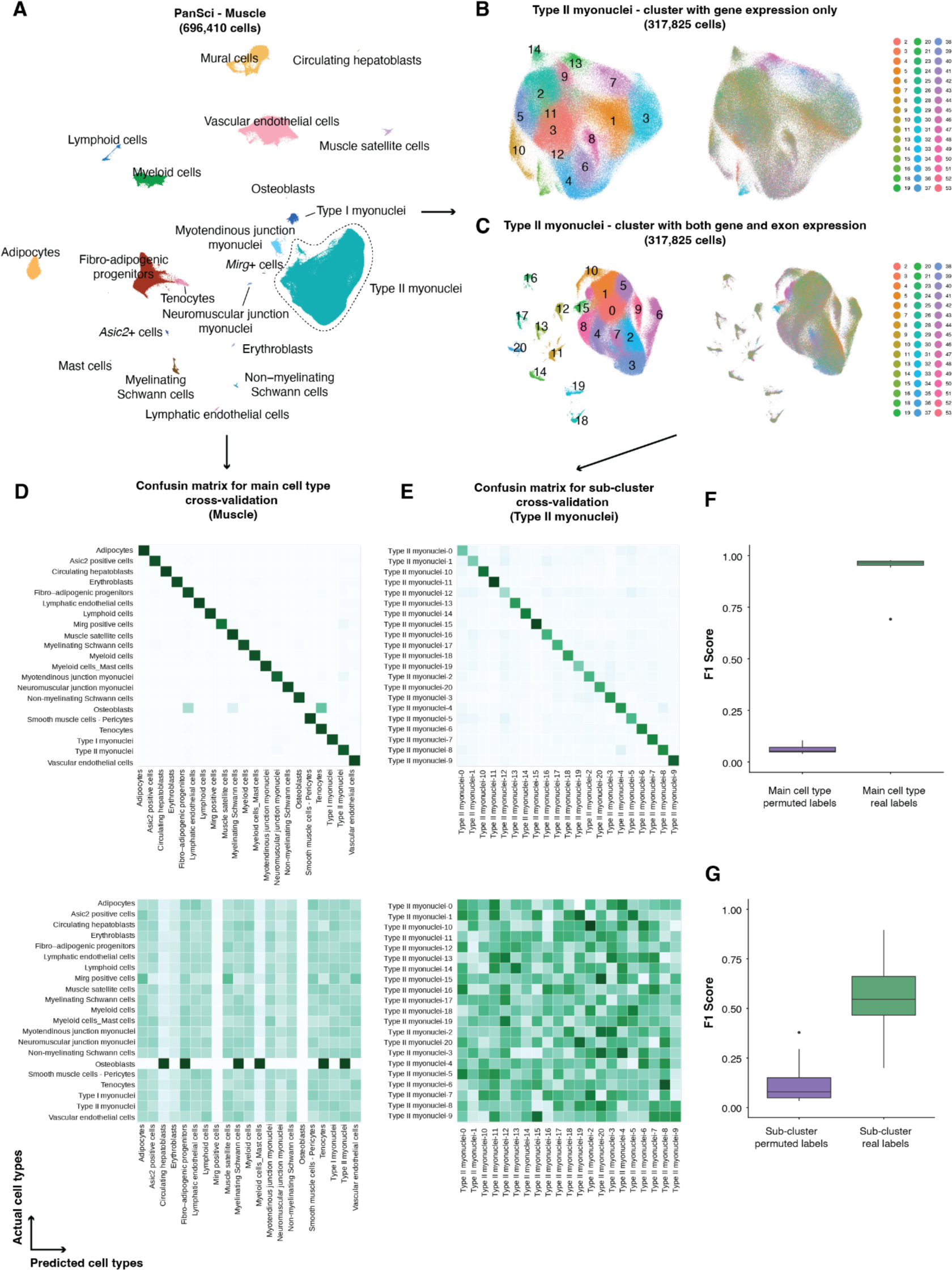
Identification and validation of main cell types and sub-clusters across organs/tissues. (**A-C**) Workflow for the identification of main cell types and sub-clusters on an organ-by-organ basis. The main cell types are initially annotated with gene markers (A). This is followed by a sub-clustering process, which utilizes combined gene and exon expression data to refine clustering resolution (B-C). (**D-G**) The cross-validation pipeline within the dataset for main cell types and subclusters is depicted. For a five-fold cross-validation, single-cell transcriptomes from these categories are input into an SVM classifier with a linear kernel (**Methods**). Confusion matrices for intra-dataset validation are generated for each organ/tissue (D) and for each main cell type (E). Additionally, specificity scores for cell annotation are determined for both main cell types (F) and subclusters (G).

**Fig. S8.**
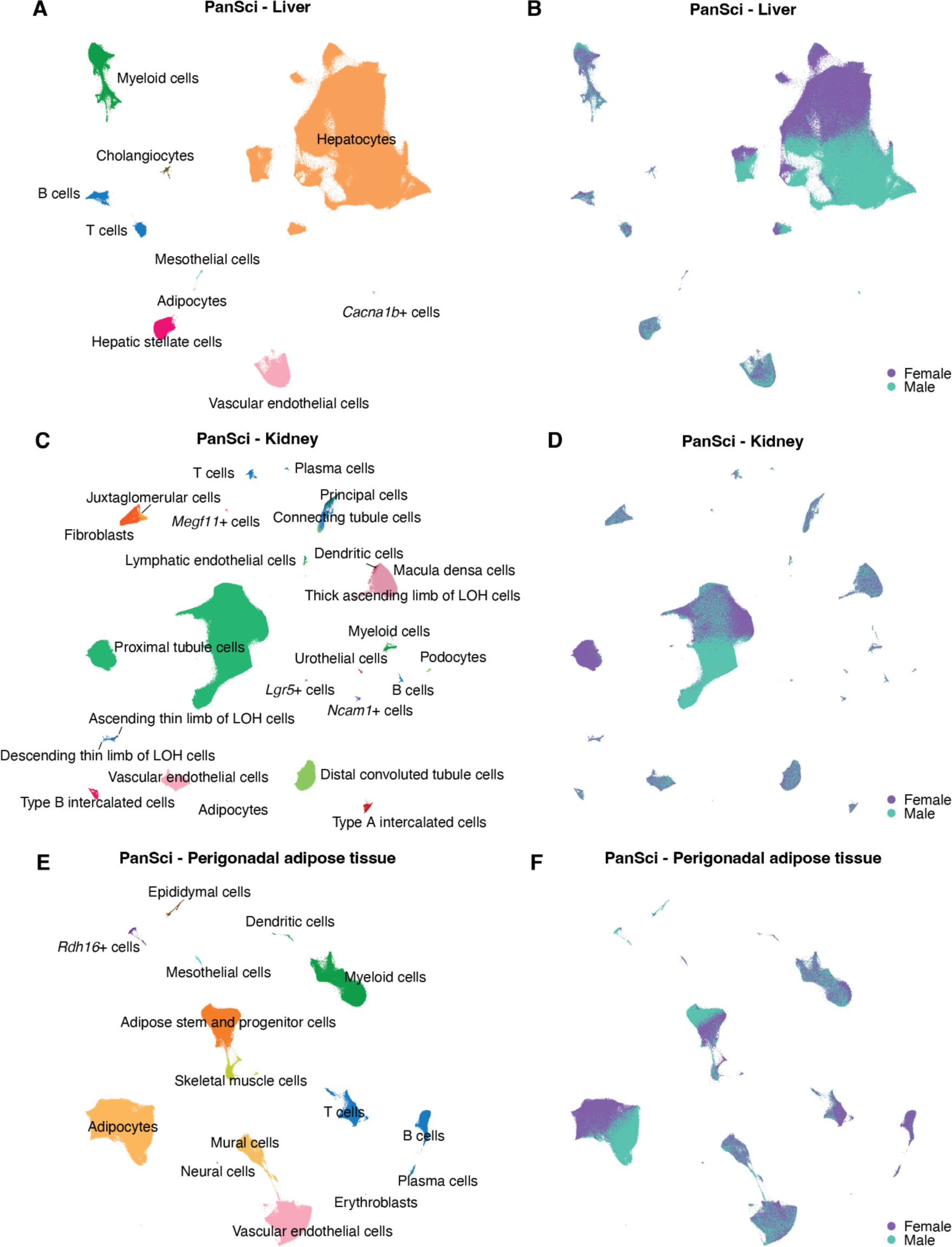
Identification of sex heterogeneity in liver, kidney, and perigonadal adipose tissue. **(A-B)** UMAP plots displaying the cellular heterogeneity in the liver, with cells color-coded by identified main cell types (A) and sexes (B). **(C-D)** UMAP plots displaying the cellular heterogeneity in the kidney, with cells color-coded by identified main cell types (C) and sexes (D). **(E-F)** UMAP plots displaying the cellular heterogeneity in the perigonadal adipose tissue, with cells color-coded by identified main cell types (E) and sexes (F).

**Fig. S9.**
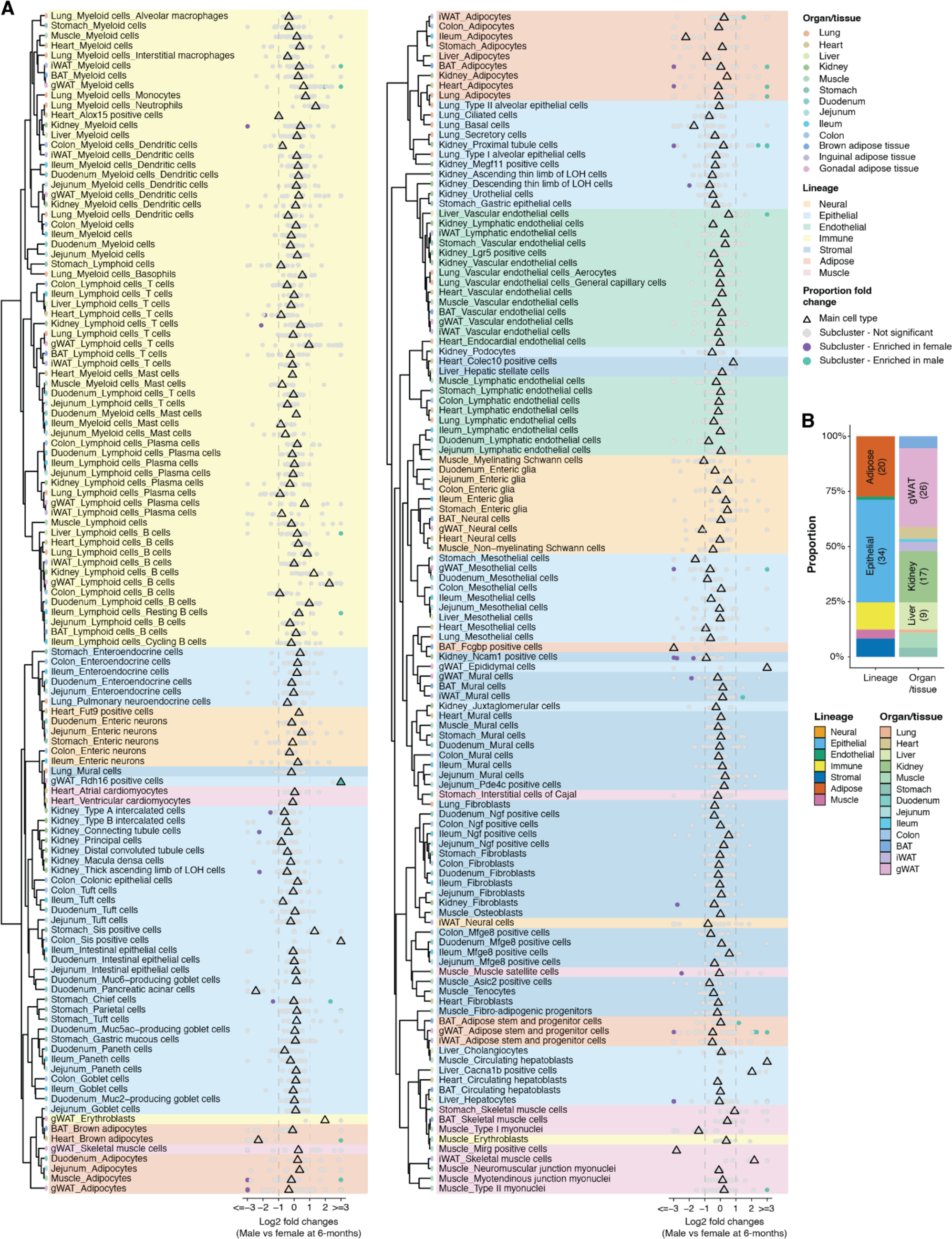
Identification of sex-specific cell types across organs/tissues. (**A**) Dot plots showing the cell-type-specific population dynamics between males and females of main cell types (triangles) and sub-clusters (dots) at 6 months old. The cell number of each main cell type and sub-cluster is normalized by the total cell numbers of each organ in respective life stages, and population dynamics is presented as the log-transformed fold changes (capped to [−3, 3]). Only cell types (both main and sub-clusters) with minimum 2-fold changes, FDR < 0.05, and consistent sex-specificity across 5 age groups are defined as sex-specific cell types. The dendrogram of each main cell type is ordered through hierarchical clustering of the correlation matrix constructed by main cell types and its top 50 principal components. (**B**) Stacked bar plots representing the proportions of sex-specific sub-clusters from different lineages and organs/tissues.

**Fig. S10.**
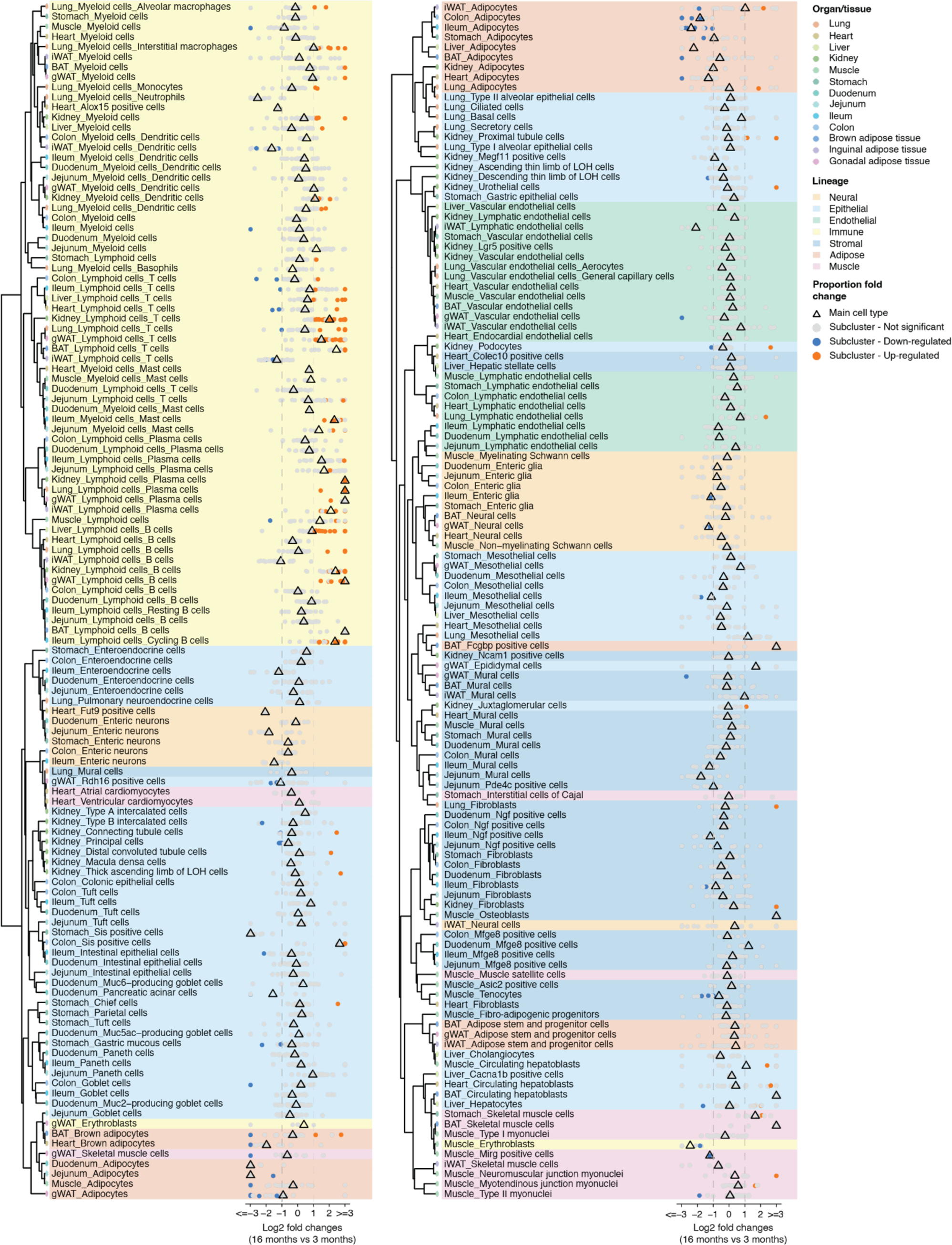
Identification of aging-associated cell population shifts across organs/tissues between 3 and 6 months. Dot plots showing the cell-type-specific population dynamics between 3 months and 16 months of main cell types (triangles) and sub-clusters (dots). The cell number of each main cell type and sub-cluster is normalized by the total cell numbers of each organ in respective life stages, and population dynamics is presented as the log-transformed fold changes (capped to [−3, 3]). Only cell types (both main and sub-clusters) with minimum 2-fold changes and FDR < 0.05 are defined as significantly changed cell types. Only significantly changed cell types consistent in both time intervals (*i.e.,* ‘3 to 16 months’ and ‘6 to 23 months’ are defined as aging-associated cell types and selected for downstream analysis. The dendrogram of each main cell type is ordered through hierarchical clustering of the correlation matrix constructed by main cell types and the top 50 principal components.

**Fig. S11:**
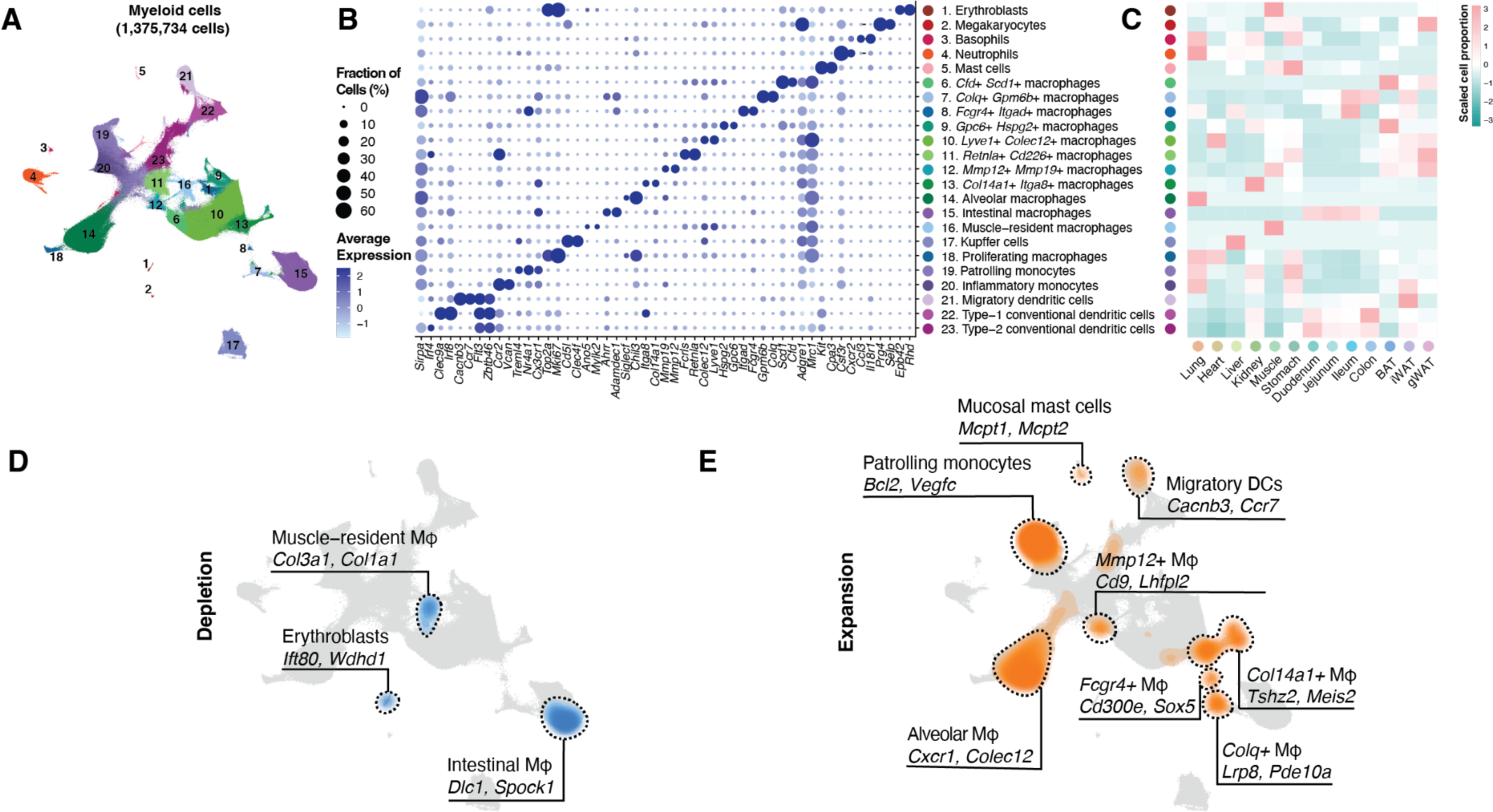
Exploration of myeloid population aging across organs/tissues. (**A**) UMAP visualizations of 1,357,734 cells from myeloid cells across organs/tissues, colored by cluster ID. (**B**) Dot plot illustrating marker gene expression for myeloid cell subtypes. The color denotes average expression values, and the dot size indicates the percentage of cells expressing these markers. (**C**) Heatmap displaying the normalized and scaled distribution of each myeloid cell subtype across organs/tissues. (**D-E**) Density plot highlighting the distribution of significantly depleted (D) and expanded (E)myeloid cells sub-clusters in aging, with their respective marker genes. (**E**) Density plot showing distribution aging-associated expansion of myeloid cells sub-clusters. Distinct marker genes are labeled for each density peak.

**Fig. S12.**
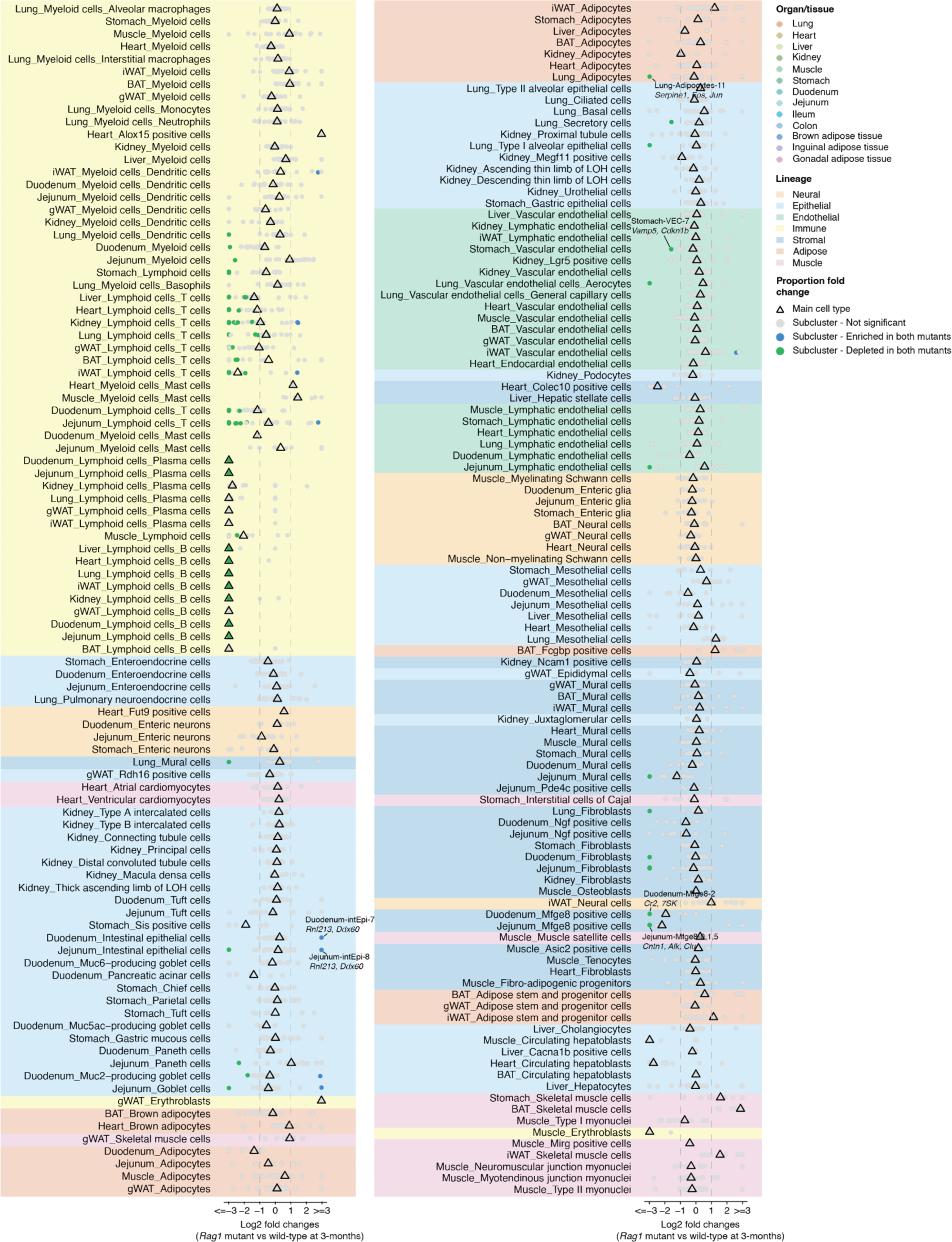
Identification of genotype-specific cell types across organs/tissues. (**A**) Dot plots showing the cell-type-specific population dynamics between Rag1 mutant and wild-type of main cell types (triangles) and sub-clusters (dots) at 3 months old. The cell number of each main cell type and sub-cluster is normalized by the total cell numbers of each organ in respective life stages, and population dynamics is presented as the log-transformed fold changes (capped to [−3, 3]). Only cell types (both main and sub-clusters) with minimum 2-fold changes, FDR < 0.05, and consistent mutant-specific enrichment/depletion are defined as genotype-specific cell types. The dendrogram of each main cell type is ordered through hierarchical clustering of the correlation matrix constructed by main cell types and its top 50 principal components.

**Fig. S13:**
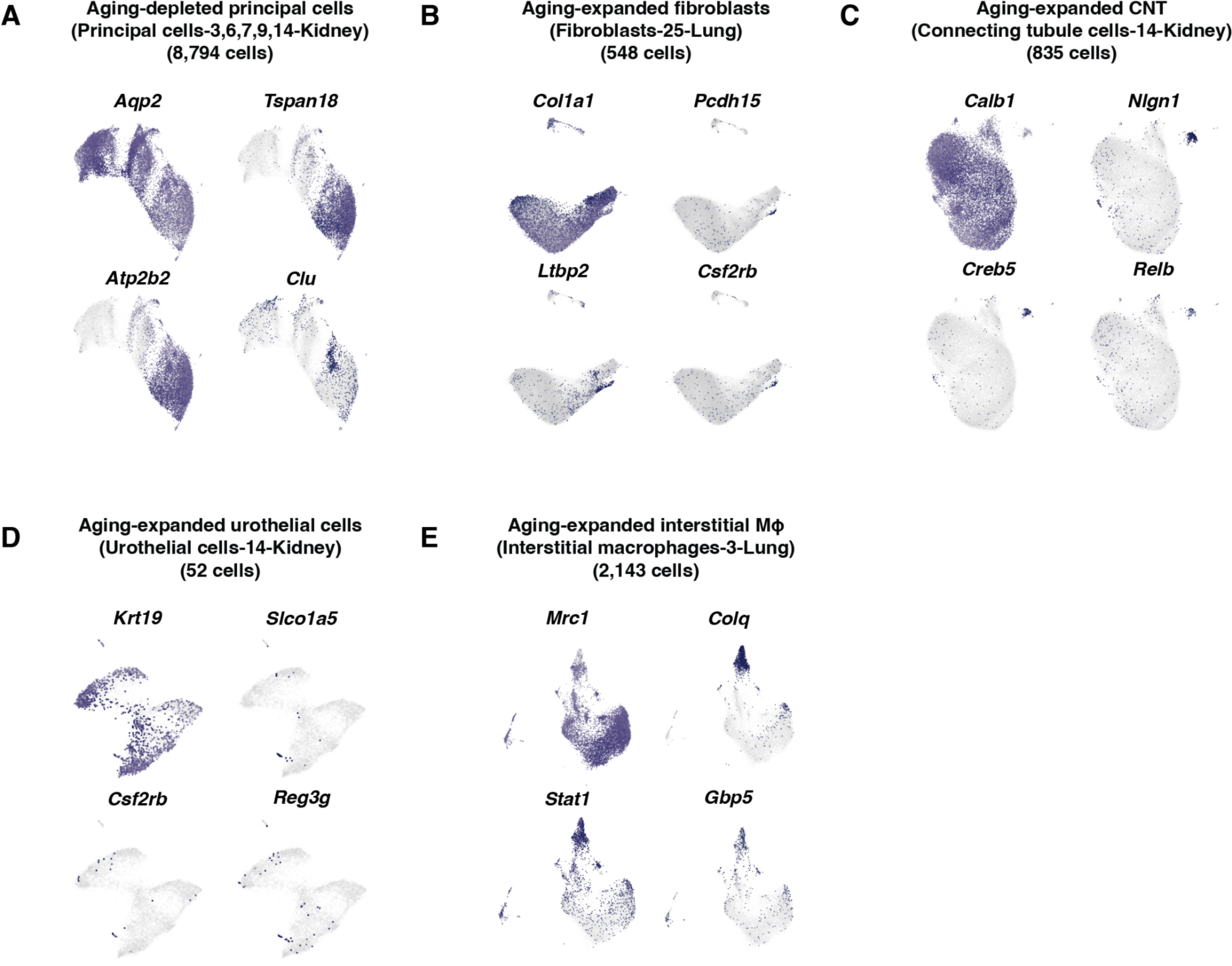
Marker gene for aging-associated sub-cluster associated with Figure 5. (A) UMAP visualization highlighting the expression of principal cell makers (*Aqp2*) and markers of aging-depleted principal cells (*Tspan18*, *Atp2b2*, and *Clu*). (B) UMAP visualization highlighting the expression of fibroblast makers (*Col1a1*) and markers of aging-expanded lung fibroblast (*Pcdh15*, *Ltbp2*, and *Csf2rb*) (K). (C) UMAP visualization highlighting the expression of CNT makers (*Calb1*) and markers of aging-expanded CNT (*Nlgn1*, *Creb5*, and *Relb*). (D) UMAP visualization highlighting the expression of urothelial cells (*Krt19*) and markers of aging-expanded urothelial cells (*Slco1a5*, *Csf2rb*, and *Reg3g*). (E) UMAP visualization highlighting the expression of interstitial macrophage makers (*Mrc1*) and markers of aging-expanded interstitial macrophage (*Colq*, *Stat1*, and *Gbp5*).

## Tables S1 to S8

**Table S1. Metadata for mouse individuals included in the study.**

**Table S2. Metadata for main cell types annotated in each organ/tissue.**

**Table S3. Differentially expressed genes for main cell types within each organ/tissue**

**Table S4. Enriched genes for sub-clusters within each main cell type**

**Table S5. List of aging-associated sub-clusters with differential abundance test results.**

**Table S6. Differentially expressed genes for T cells and innate lymphoid cells subtypes**

**Table S7. Differentially expressed genes for B cells and plasma cells subtypes.**

**Table S8. Differentially expressed genes for myeloid cells subtypes.**

